# Distinct Classes of Gut Bacterial Molybdenum-Dependent Enzymes Produce Urolithins

**DOI:** 10.1101/2025.01.17.632430

**Authors:** Minwoo Bae, Xueyang Dong, Julian Avila-Pacheco, Fechi Inyama, Vayu Hill-Maini, Clary Clish, Emily P. Balskus

## Abstract

Urolithin A is an anti-aging and anti-inflammatory gut bacterial metabolite derived from ellagic acid (EA), a polyphenol abundant in berries and nuts. The conversion of EA to urolithin A involves multiple chemically challenging phenol dehydroxylation steps that produce urolithins with varying bioactivities. Despite their biological and chemical significance, the bacterial enzymes responsible for urolithin production remain unidentified. Here, we use differential gene expression analysis, anaerobic protein expression, and enzyme assays to identify members of two distinct molybdenum enzyme families (the DMSO reductase family and the xanthine oxidase family) capable of regioselective dehydroxylation and urolithin generation. These two enzyme families have distinct substrate requirements, suggesting they employ different catalytic mechanisms for phenol dehydroxylation. Multi-omics analysis of a human cohort uncovers decreased levels of urolithin A and genes encoding urolithin A-producing enzymes in patients with inflammatory bowel disease (IBD), implying reduced health effects of EA consumption in this setting. Together, this study elucidates the molecular basis of urolithin production, expands the known enzymatic repertoire of the human gut microbiome, and suggests a potential link between gut bacterial urolithin production and host inflammation.

**Significance statement:** The human gut microbiome modulates the health effects of dietary compounds by modifying their chemical structures. For example, gut microbes extensively metabolize polyphenols, a group of diverse plant-derived compounds associated with positive health outcomes. Urolithin A, a gut bacterial metabolite derived from a polyphenol abundant in berries and nuts, exhibits potent anti-inflammatory activity. However, the gut bacterial enzymes involved in its production remain largely unknown. Here, we report urolithin-producing gut bacterial enzymes, including four phenol dehydroxylases from two distinct molybdenum-dependent enzyme families. Analyzing human gut microbiomes suggests a potential link between genes encoding these enzymes and host inflammation. Together, our findings fully map urolithin A production to enzymes, increasing our understanding of how the gut microbiome can alter the impacts of diet on human health.

## Introduction

EA is a polyphenol derived from ellagitannins abundant in foods such as berries, nuts, and pomegranates^1^. Consumption of EA-rich foods is associated with positive health outcomes such as improved cardiovascular health and decreased mortality^2–5^. These health benefits have partly been attributed to the urolithins, a group of EA-derived metabolites produced by gut bacteria^6^. Urolithin A, the urolithin whose bioactivity has been most extensively studied, inhibits pro-inflammatory cytokine production and restores mitophagy in aged cells^7–11^. In model organisms, the activities of urolithin A have been linked to reduced inflammation^12–14^, activation of anti-tumor immunity^15^, and protection against age-related conditions such as repressed stem cell function^16^, muscle dysfunctions^7,17,8^, neurological disorders^18–20^, and metabolic diseases^21–23^ (**Figure 1A**). Recent clinical studies demonstrated that urolithin A administration improved muscle function in elderly and middle aged human subjects^24–26^, and urolithin A is currently marketed as a dietary supplement in the US. Importantly, fecal urolithin A levels vary widely across individuals even when EA consumption is controlled, presumably due to varied gut microbial metabolic capabilities^27–29^. Despite the biological importance of gut bacterial urolithin A production, the genes and enzymes involved in this metabolism remain to be uncovered.

**Figure 1.**
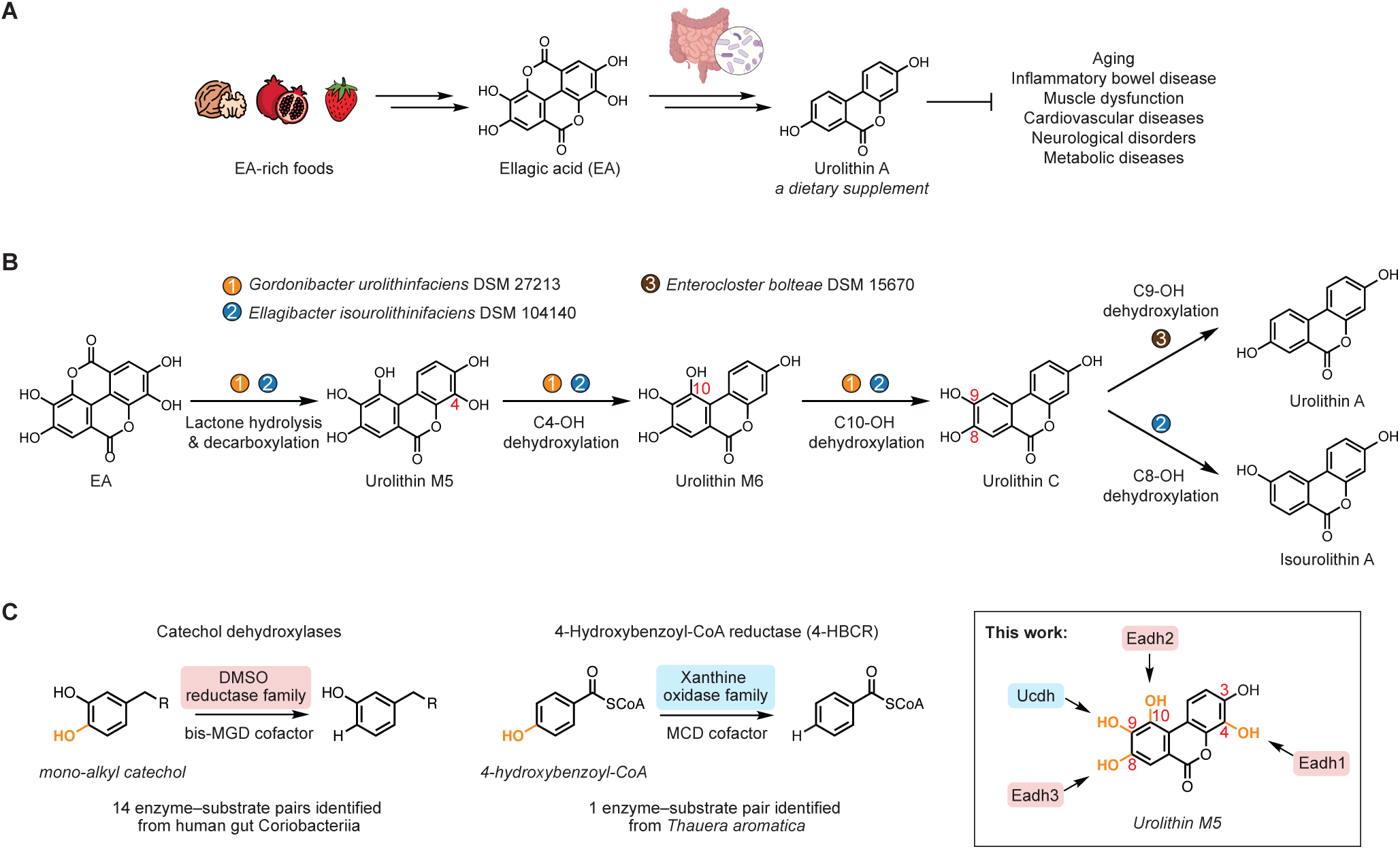
Unknown gut bacterial enzymes produce urolithin A. **(A)** The beneficial health effects of urolithin A are derived from gut microbial metabolism of the dietary polyphenol ellagic acid (EA). **(B)** Major metabolic pathways for EA by previously reported gut bacterial species. The pathways involve urolithins with different hydroxylation patterns. **(C)** Comparison between phenol dehydroxylations performed by known enzymes (catechol dehydroxylases and 4-HBCR) and dehydroxylations involved in EA metabolism. Phenol groups of urolithin M5 subject to dehydroxylation are highlighted in orange.

Intriguingly, the conversion of EA to urolithin A involves a series of phenol dehydroxylations that yields urolithins with different hydroxylation patterns. Recent studies have shown that the human gut Coriobacteriia species *Gordonibacter urolithinfaciens* converts EA to urolithin C, which is further metabolized to urolithin A by *Enterocloster bolteae*, a gut Clostridial species (**Figure 1B**)^30,31^. These two species remove three hydroxyl groups of urolithin M5 at C4, C10, and C9 to produce urolithin M6, urolithin C, and urolithin A in sequence. Alternatively, the gut Coriobacteriia species *Ellagibacter isourolithinifaciens* produces a regioisomer of urolithin A called isourolithin A via dehydroxylation of urolithin C at C8 (**Figure 1B**)^32^. Importantly, despite the subtle changes in their structures, urolithins with different hydroxylation patterns exhibit different bioactivity *in vitro*^33–35^. For example, urolithin A, but not isourolithin A, significantly reduced triglyceride accumulation in human adipocytes^33^. Dietary intervention studies have revealed that individuals who metabolize EA to urolithin A have improved health outcomes, including lower rates of obesity and lower levels of cardiovascular disease biomarkers compared to a group of people who produce isourolithin A and its congener urolithin B^36–38^. Therefore, the regioselectivity of urolithin dehydroxylation may affect the health consequence of EA consumption.

Phenol dehydroxylation is a transformation of high chemical interest as it is a key reaction in lignin valorization ^39,40^. However, current methods are energy-intensive, requiring high temperature and high H2 partial pressure to activate the stable aromatic C–O bond^41^. Interestingly, this transformation is central to anaerobic bacterial metabolism of aromatic compounds and has been linked to two types of molybdenum-dependent enzymes (**Figure 1C**). Notably, catechol dehydroxylases, members of the dimethyl sulfoxide (DMSO) reductase family, catalyze the dehydroxylation of a catechol motif (i.e., 1,2-dihydroxylated benzene) using a bis-molybdopterin guanidine dinucleotide (bis-MGD) cofactor^42–46^. Primarily encoded in the genomes of gut Coriobacteriia, characterized catechol dehydroxylases process polyphenols with high substrate specificity^42–44,47,48^ and have been shown to support anaerobic respiration^44,47,49^. Another phenol dehydroxylase, 4-hydroxybenzoyl-CoA reductase (4-HBCR), removes the hydroxyl group of 4-hydroxybenzoyl-CoA, a key step in anaerobic catabolism of aromatic compounds^50^. Found in the sewage-derived bacterium *Thauera aromatica*^51^, 4-HBCR is a member of the xanthine oxidase family and carries a molybdenum cytosine dinucleotide (MCD) cofactor^52^. The serial dehydroxylations observed in EA bioactivation, which occur on a highly oxygenated aromatic scaffold, offer an exciting opportunity for further discovery of phenol dehydroxylases as they are distinct from these previously reported transformations.

Here, we elucidate the gut bacterial enzymes involved in the complete conversion of EA to urolithin A and the competing pathways to isourolithin A and urolithin B. Using differential gene expression analysis, protein expression, and enzyme activity assays, we identify four urolithin dehydroxylases: three catechol dehydroxylases (Eadh1, Eadh2, and Eadh3) from *Gordonibacter* strains and *E. isourolithinifaciens* and a xanthine oxidase family enzyme (Ucdh) from *E. bolteae* (**Figure 1C**). By examining substrate scope and enzyme kinetics, we reveal that these enzymes target individual hydroxyl groups on the urolithins and exhibit substrate specificity that explains the sequence of this metabolic pathway. Finally, we find that both Coriobacteriia EA-metabolizing gene abundance and urolithin A levels are depleted in IBD patients, suggesting a potential route through which inflammation alters production of the bioactive metabolite urolithin A. Altogether, this work elucidates the molecular basis of gut bacterial urolithin production and lays the groundwork for future efforts to understand polyphenol dehydroxylation and its relevance for human health.

## Results

### Identification of a Coriobacteriia gene cluster responsible for EA metabolism

We hypothesized that *Gordonibacter* and *Ellagibacter* species likely employ catechol dehydroxylases for urolithin metabolism, given that strains of these species typically encode over 30 uncharacterized members of this enzyme family^47^. Since catechol dehydroxylases are also widespread across other gut Coriobacteriia, we screened a library of Coriobacteriia strains for EA metabolism in liquid culture using LC–MS/MS quantification. Consistent with previous findings^30,32^, urolithin C was produced by *G. urolithinfaciens* and *Gordonibacter pamelaeae*, while isourolithin A was generated by *E. isourolithinifaciens* (**Figure S1A**). In cultures of *G. urolithinfaciens* and *G. pamelaeae*, the intermediates urolithin M5 and urolithin M6 were also detected. Additionally, we found another human gut *Gordonibacter* strain, *Gordonibacter* sp. (*Gs*) 28C, completely converted EA to urolithin C. All other Coriobacteriia strains we tested were inactive (only EA was recovered), and none of the strains produced urolithin A. We then investigated whether this metabolic activity is induced by EA, as transcription of genes encoding catechol dehydroxylases is typically induced by their cognate substrates. The cell lysate of *Gs* 28C converted EA to urolithin C when the culture was grown with EA, whereas lysate from vehicle-treated culture showed no activity (**Figure S1B**). This result indicated that the enzymes responsible for EA metabolism are expressed in response to the substrate, similar to other catechol dehydroxylases^42–44,53^.

To identify putative EA-metabolizing genes, we performed a differential gene expression analysis of *Gs* 28C treated with either EA or vehicle, a strategy that has been previously used to link catechol substrates to their dehydroxylases^42–44^. We identified thirteen genes that were upregulated by more than 1,000-fold in response to EA compared to vehicle (**Figure 2A**). These genes mapped onto a single gene cluster that encoded two putative molybdopterin-dependent oxidoreductases (Eadh1 and Eadh2) and additional proteins related to EA metabolism. A protein phylogenetic analysis placed these two enzymes in the catechol dehydroxylase clade alongside known catechol dehydroxylases from *Gordonibacter* strains (**Figure S1C**). Like other *Gordonibacter*-type catechol dehydroxylases^44^, the genes encoding Eadh1 and Eadh2 are colocalized with genes encoding 4Fe-4S binding proteins (**Figure 2A**). Considering that EA metabolism by *Gs* 28C involves two dehydroxylation steps, we reasoned that Eadh1 and Eadh2 may each catalyze one step. By performing a sequence-based homology search with Eadh1 and Eadh2 as queries using Basic Local Alignment Search Tool (BLAST)^54^, we identified homologous gene clusters in all known EA metabolizers as well as as-yet-uncultivated Coriobacteriia from human and other animal gut microbiomes (**Figure 2B** and **Figure S1D**). Interestingly, the gene cluster from *E. isourolithinifaciens*, which metabolizes EA to isourolithin A using one additional dehydroxylation step^32^, encoded an additional uncharacterized molybdopterin-dependent oxidoreductase (Eadh3) that is also found within the catechol dehydroxylase clade (**Figure 2B** and **Figure S1C**). We hypothesized that this enzyme processes urolithin C to isourolithin A.

**Figure 2.**
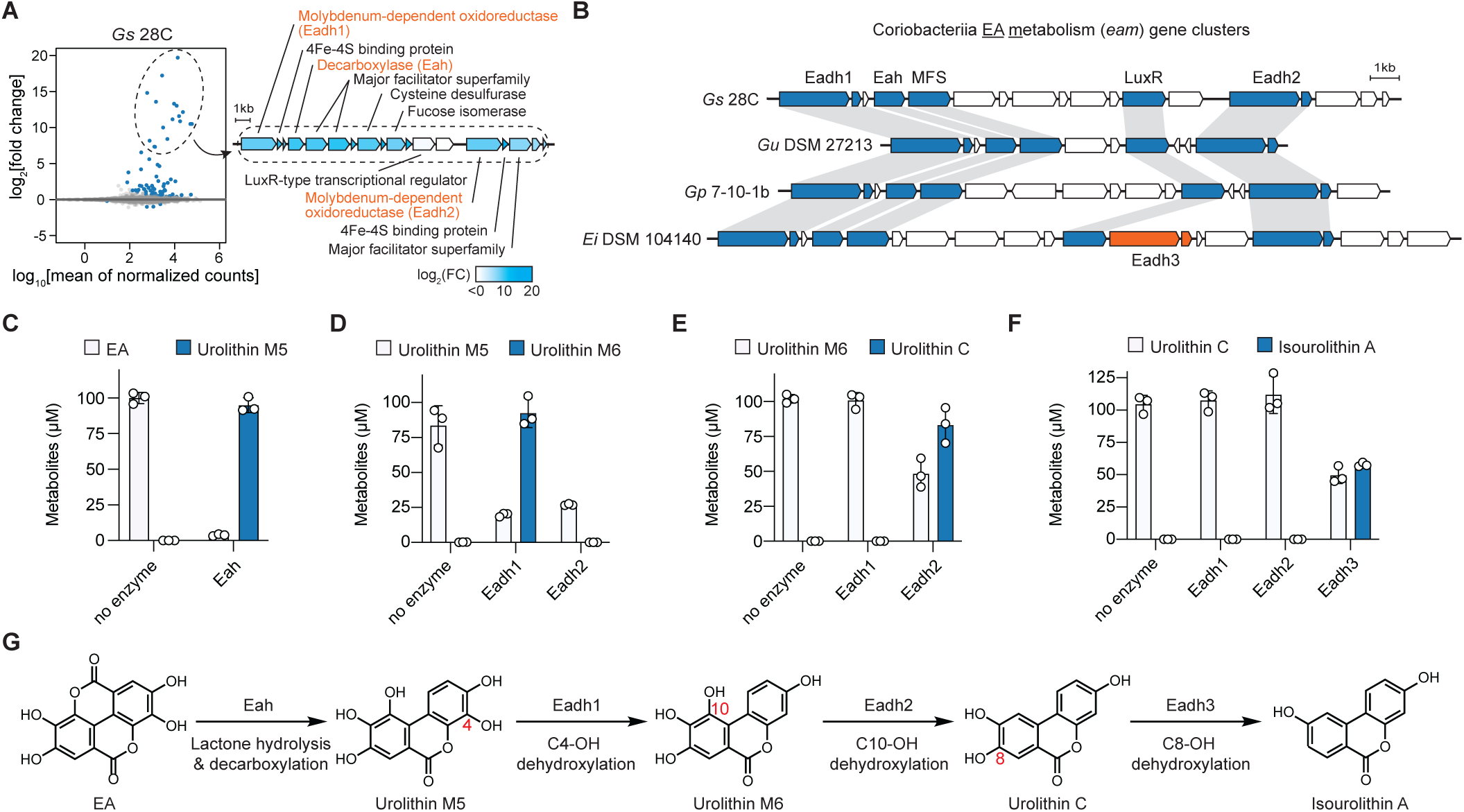
Identification of a Coriobacteriia gene cluster responsible for EA metabolism. **(A)** Left panel: differential gene expression analysis of *Gs* 28C cultured in media supplemented with 100 µM EA or vehicle (n=3 biological replicates). Genes with adjusted *p* value < 0.01 are represented by navy dots. Right panel: functional annotations for proteins encoded by the gene cluster induced by EA. **(B)** Conservation of the EA metabolism (*eam*) gene cluster among EA-metabolizing Coriobacteriia strains. The conserved genes are colored navy. Genes encoding an additional catechol dehydroxylase homolog (Eadh3) in *E. isourolithinifaciens* are colored orange. **(C)** Transformation of EA to urolithin M5 by Eah. 1 µM Eah was incubated with 100 µM EA at rt for 20 h in a pH 7.0 buffer (50 mM HEPES and 250 mM NaCl) under anaerobic conditions. **(D)** Metabolism of urolithin M5 by Eadh1 and Eadh2. **(E)** Metabolism of urolithin M6 by Eadh1 and Eadh2. **(F)** Transformation of urolithin C by Eadh1, Eadh2, and Eadh3. **(D–F)** 100 nM enzyme was incubated with 100 µM substrate, 200 µM methyl viologen, and 100 µM sodium dithionite at rt for 20 h in a pH 7.0 buffer (20 mM MOPS and 300 mM NaCl) under anaerobic conditions. **(C–F)** Data represented as mean ± SD with n = 3 replicates. **(G)** Assignment of enzymes in the complete EA metabolic pathway by *Gordonibacter* and *Ellagibacter* strains. *Gs, Gordonibacter species*; *Gu, Gordonibacter urolithinfaciens*; *Gp, Gordonibacter pamelaeae*; *Ei, Ellagibacter isourolithinifaciens*; MFS, major facilitator superfamily.

In addition to the putative catechol dehydroxylases, the identified gene clusters all contained genes encoding a decarboxylase (Eah), MFS transporters, and a LuxR-type transcriptional regulator. MFS transporters and LuxR-type transcriptional regulators are often found in the genomic neighborhood of catechol dehydroxylases. MFS transporters are thought to transport substrates, and LuxR-type regulators have been shown to regulate transcription of genes encoding other catechol dehydroxylases^44,53^. Annotated as a metal-dependent decarboxylase (IPR032465) by InterPro 98.0^55^, Eah is a candidate for the initial lactone ring opening and decarboxylation of EA to urolithin M5. We heterologously expressed Eah from *Gs* 28C in *Escherichia coli* BL21 (DE3) and confirmed that the purified protein converted EA to urolithin M5 (**Figure 2C** and **Figure S1E**). This suggested that the gene cluster identified by differential gene expression analysis was likely responsible for EA metabolism, and we named this gene cluster as the Coriobacteriia <u>EA m</u>etabolism (*eam*) gene cluster.

Next, we sought to characterize the catechol dehydroxylase homologs (Eadh1/2/3) associated with this gene cluster. Using a recently developed cumate-inducible expression system for catechol dehydroxylases^53^, we anaerobically expressed and purified recombinant Eadh1 and Eadh2 from *G. urolithinfaciens,* their native host (**Figure S1F**). Each purified enzyme was incubated with either urolithin M5 or urolithin M6 along with methyl viologen and sodium dithionite under anaerobic conditions, and urolithin substrates and products were quantified using LC– MS/MS. Consistent with our hypothesis, distinct dehydroxylation steps were catalyzed by Eadh1 (C4-OH dehydroxylation of urolithin M5 to urolithin M6) (**Figure 2D**) and Eadh2 (C10-OH dehydroxylation of urolithin M6 to urolithin C) (**Figure 2E**). Similarly, Eadh3 from *E. isourolithinifaciens* was expressed in *G. urolithinfaciens* and purified for enzyme assays (**Figure S1F**). As predicted, Eadh3 converted urolithin C to isourolithin A via C8-OH dehydroxylation (**Figure 2F**). Altogether, the activities of Eah, Eadh1, Eadh2, and Eadh3 account for the complete EA metabolic pathway in *Gordonibacter* and *Ellagibacter* strains (**Figure 2G**).

### Catechol dehydroxylase substrate specificity and kinetics explain urolithin production

Surprisingly, we noticed that Eadh2 also consumed urolithin M5 but did not produce urolithin M6 (**Figure 2D**). Instead, it generated a new mass feature that had the same mass as urolithin M6 but eluted at a different retention time on LC–MS/MS (**Figure 3A**). Considering that Eadh2 dehydroxylated the C10-OH of urolithin M6, we reasoned this new product could be urolithin D, the C10-OH dehydroxylated metabolite of urolithin M5^56^. Indeed, the retention time of the product produced by Eadh2 from urolithin M5 matched that of a urolithin D synthetic standard (**Figure 3A**). These results showed that Eadh2 dehydroxylates the C10-OH of both urolithin M5 and urolithin M6 substrates, which differ only by a hydroxyl on C4. Likewise, Eadh1, the enzyme removing C4-OH of urolithin M5, can accept urolithin D as a substrate to produce urolithin C via dehydroxylation at C4 (**Figure S2A**). Eadh3 also demonstrated activity towards multiple C8-OH containing compounds other than urolithin C, but product identities could not be verified due to lack of authentic standards (**Figure S2B**). Importantly, dehydroxylation by these enzymes required the hydroxyl group being removed to be part of a catechol. For example, urolithin E, which has its C4-OH incorporated into a catechol, but not its C10 and C8-OHs, was accepted as a substrate only by the C4-OH dehydroxylating Eadh1 (**Figure 3B**). This requirement is consistent with previous studies of known catechol dehydroxylases, which revealed that a catechol is essential for activity^44^. Combined, these results suggest that Eadh1, Eadh2, and Eadh3 regioselectively remove catecholic C4, C10, and C8-OHs of urolithins, respectively, with substrate promiscuity in terms of non-participating hydroxyl substituents.

**Figure 3.**
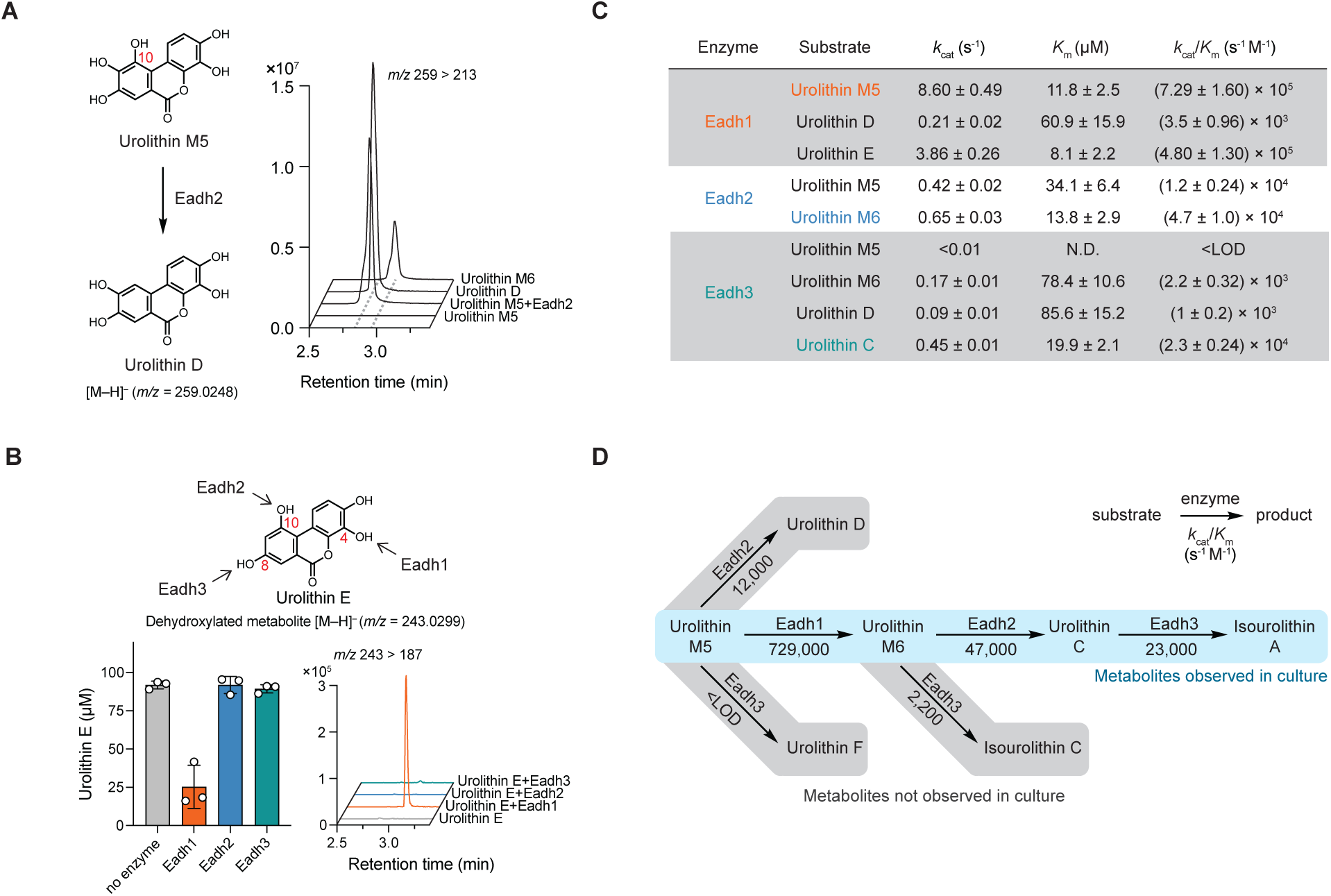
Catechol dehydroxylase substrate specificity and kinetics explain urolithin production. **(A)** LC–MS/MS traces of urolithin M5 metabolism by Eadh2. **(B)** Urolithin E dehydroxylation by Eadh1, Eadh2, and Eadh3 (n=3). **(A–B)** 100 nM enzyme was incubated with 100 µM substrate, 200 µM methyl viologen, and 100 µM sodium dithionite at rt for 20 h in pH 7.0 buffer (20 mM MOPS and 300 mM NaCl) under anaerobic conditions. **(C)** Kinetic parameters of Eadh1, Eadh2, and Eadh3 using initial rates measured in the first 60 s. Data are mean ± SD (n=3). Fitted parameters are mean ± SE (n=3) as derived from nonlinear curve fitting to the Michaelis–Menten equation. The reactions were performed in pH 7.0 buffer (20 mM MOPS and 300 mM NaCl) containing varying concentrations of enzymes and substrates, 200 µM methyl viologen, and 100 µM sodium dithionite at 30 °C. **(D)** A scheme of possible metabolic routes for urolithins by Eadh1, Eadh2, and Eadh3 and the catalytic efficiency (*k*cat/*K*m) of each step. LOD, limit of detection; N.D., not determined.

The promiscuity of these catechol dehydroxylases implied possible alternative EA metabolic pathways. For example, urolithin M5 could be dehydroxylated *in vitro* by Eadh1 and Eadh2 to urolithin M6 and urolithin D, respectively. However, urolithin D has not been detected in *Gordonibacter* or *Ellagibacter* liquid cultures incubated with EA, for which urolithin M6 has been proposed as the major intermediate^30,32^. We reasoned that the specific intermediates observed in culture are likely determined by the substrate preferences of Eadh enzymes. To test this hypothesis, we performed enzyme kinetics experiments for each dehydroxylase–substrate pair using a previously reported continuous absorbance-based assay^47^. All three enzymes showed varying kinetic parameters towards different urolithin substrates (**Figure 3C** and **Figure S2C**). Comparing the kinetic efficiency (*k*cat/*K*m) for different substrates revealed that urolithin M5, urolithin M6, and urolithin C are the preferred substrates of Eadh1, Eadh2, and Eadh3, respectively. Importantly, these preferences are consistent with the sequence of intermediates observed in culture. For example, the predominant production of urolithin M6 from urolithin M5 by *Gordonibacter* and *Ellagibacter* strains can be attributed to the significantly higher kinetic efficiency of Eadh1 towards urolithin M5 (729,000 s^-1^ M^-1^) compared to the other two enzymes (12,000 s^-1^ M^-1^ by Eadh2 and <limit of detection by Eadh3) (**Figure 3D**). Similarly, urolithin M6 was metabolized more readily by Eadh2 than by Eadh3, which aligns with the observed formation of urolithin C as a precursor of isourolithin A. Altogether, these enzyme kinetics experiments revealed the native substrates for each dehydroxylase and explain the sequence of EA metabolism observed in culture.

### A previously unappreciated xanthine oxidase homolog catalyzes dehydroxylation of urolithin C to urolithin A

Next, we sought to identify the *E. bolteae* enzyme responsible for the C9-OH dehydroxylation of urolithin C to urolithin A. To date, phenol dehydroxylations within the human gut have been exclusively attributed to catechol dehydroxylases from Coriobacteriia^42–44,47,48^. However, *E. bolteae,* a Clostridial species, encodes two DMSO reductase family enzymes (IPR006657) in its genome, both of which share <22% amino acid identity to any characterized catechol dehydroxylases, suggesting that this step may not be catalyzed by this type of enzyme. Notably, unlike the hydroxyl groups removed by characterized catechol dehydroxylases, the C9-OH of urolithin C is positioned *para* to an electron-withdrawing ester group. This electronically resembles 4-hydroxybenzoyl-CoA, the substrate of the xanthine oxidase family enzyme 4-HBCR (**Figure 1C**), which contains a phenol group *para* to a thioester, a configuration proposed to stabilize catalytic intermediates^57,58^. This analysis suggested that a protein related to 4-HBCR may be involved in the C9-OH dehydroxylation.

The *E. bolteae* genome encodes 13 proteins annotated as xanthine oxidases (IPR016208) whose amino acid sequence identity to 4-HBCR ranges from 23% to 34%. To determine if differential gene expression analysis could pinpoint candidate enzymes, we tested whether the urolithin C dehydroxylation activity is induced by substrate. The *E. bolteae* cell suspension from a culture grown with urolithin C showed activity, whereas the suspension from a vehicle-treated culture lacked activity (**Figure 4A**). Thus, we applied a differential gene expression analysis to *E. bolteae* treated with urolithin C or vehicle and identified two gene clusters (cluster I and cluster II) upregulated in the presence of the substrate (**Figure 4B**). Cluster I contained functionally diverse protein-encoding genes, including transcriptional regulators, a methyltransferase, and three subunits of a xanthine oxidase family enzyme (UcdhABC): a catalytic subunit harboring the MCD cofactor (UcdhC), a medium subunit with a flavin adenine dinucleotide (FAD) cofactor (UcdhA), and a small subunit with two 2Fe-2S clusters (UcdhB). Notably, 4-HBCR was the closest characterized member of the xanthine oxidase family to Ucdh (32.7% amino acid identity, 49.2% amino acid similarity) (**Figure 4C**). Cluster II encoded proteins involved in molybdate transport and molybdopterin cofactor biosynthesis, maturation, and insertion, including XdhC, a chaperone that aids MCD maturation and insertion for xanthine oxidase family enzymes^59^. Given these results, we hypothesized a role for Ucdh in urolithin C dehydroxylation.

**Figure 4.**
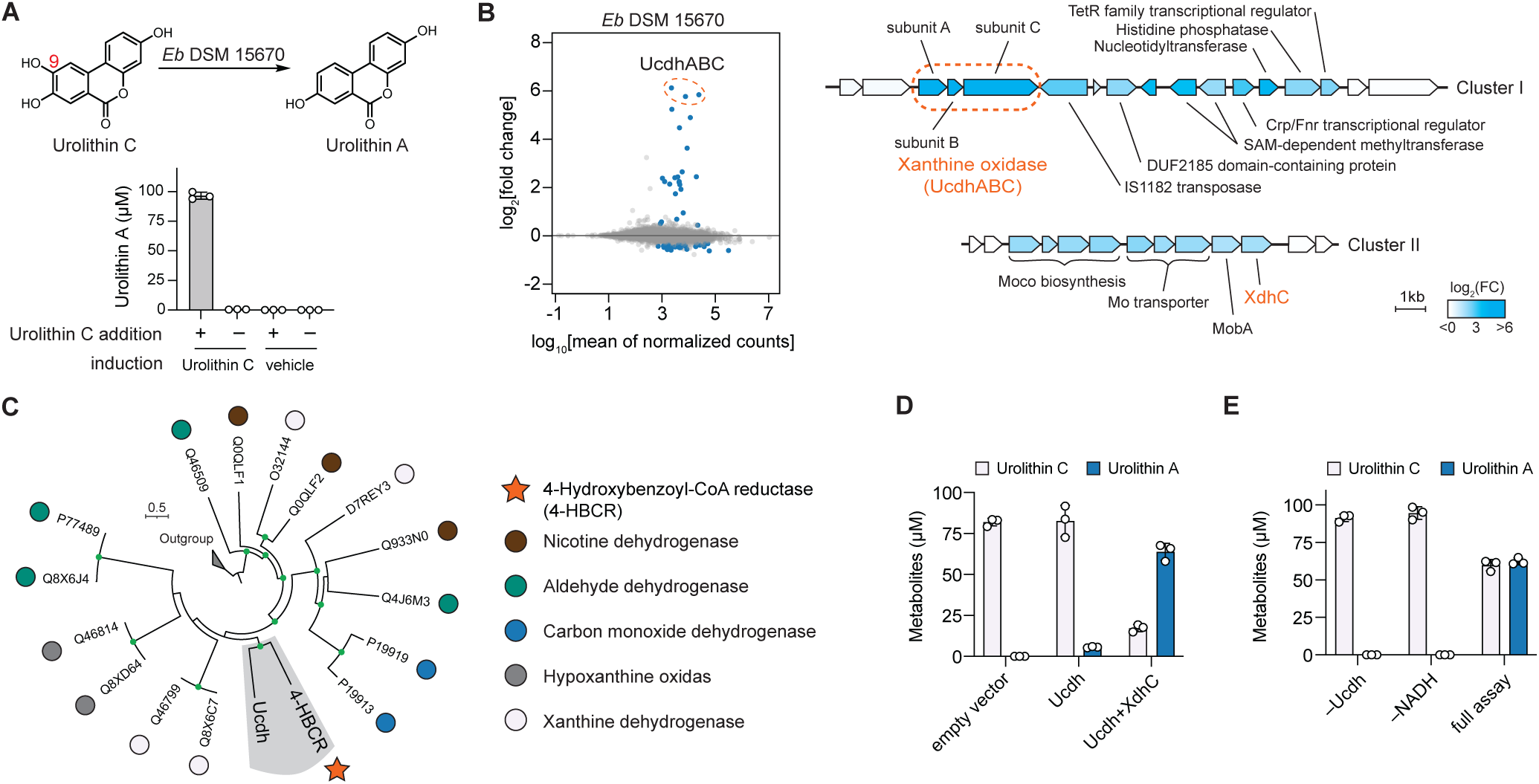
A previously unappreciated xanthine oxidase homolog catalyzes dehydroxylation of urolithin C to urolithin A. **(A)** Cell suspensions of *Eb* DSM 15670 induced by 100 µM urolithin C or vehicle were incubated with 100 µM urolithin C or vehicle for 6 h. **(B)** (left panel) Differential gene expression in *Eb* DSM 15670 induced by 100 µM urolithin C or vehicle. Genes with adjusted *p* values < 0.05 were colored in navy. (right panel) Gene annotations of the two gene clusters upregulated in response to urolithin C treatment. **(C)** Phylogenetic tree of biochemically characterized bacterial xanthine oxidase enzymes and Ucdh. Nodes with bootstrap value > 0.7 are shown in green. The outgroup represents eukaryotic xanthine oxidase enzymes. **(D)** Metabolism of urolithin C by *E. coli* TP1000 cell suspension expressing Ucdh alone or Ucdh and XdhC under anaerobic conditions (n=3 biological replicates). **(E)** Dehydroxylation of urolithin C by purified Ucdh (n=3). The full assay was performed anaerobically by adding 2 µM Ucdh to a pH 7.0 buffer (20 mM MOPS, 300 mM NaCl) containing 100 µM urolithin C and 1 mM NADH. *Eb*, *Enterocloster bolteae*

To confirm its enzymatic activity, we sought to obtain active Ucdh by expressing the protein under anaerobic conditions given the oxygen sensitivity of 4-HBCR^60^. Our initial attempt to express Ucdh in *E. coli* BL21 (DE3) yielded inactive protein, presumably due to challenges with MCD maturation and insertion. To overcome this issue, we used *E. coli* TP1000^61^, an engineered host which overproduces the MCD cofactor. TP1000 cell suspensions expressing Ucdh converted urolithin C to urolithin A, and co-expression with the putative chaperone XdhC from cluster II further increased urolithin A production (**Figure 4D**). The Ucdh purified from this expression system converted urolithin C to urolithin A with the addition of NADH as an electron donor (**Figure 4E** and **Figure S3A**). Ucdh also regioselectively dehydroxylated other C9-OH containing urolithins at this position (**Figure S3B**), including isourolithin A whose C9-OH is not part of a catechol, further illustrating that its activity is distinct from that of catechol dehydroxylases. In summary, we have identified the xanthine oxidase enzyme Ucdh as a urolithin C9-OH dehydroxylase.

To understand the distribution of Ucdh homologs in sequenced bacteria, we searched the National Center for Biotechnology Information (NCBI) RefSeq Select protein database using BLAST. High homology hits (>90% amino acid identity) were exclusively identified in the genomes of *E. bolteae* and its relatives like the human gut bacteria *Enterocloster asparagiformis* and *Enterocloster hominis* (formerly known as *Lachnoclostridium pacaense*^62^) (**Table S1** and **Figure S3D**)^63^. *E. asparagiformis* dehydroxylated urolithin C in culture, suggesting that the Ucdh homologs in these strains are likely active (**Figure S3F**). All other hits shared less than 60% amino acid identity with Ucdh (**Figure S3D)**. Notably, most species that encode these low homology hits are anaerobes isolated from environments like sludge, marine sediment, and wastewater (**Table S1**). Whether these environmental Ucdh homologs also dehydroxylate urolithin C is currently unknown. Alternatively, these enzyme may catalyze other reductive transformations related to anaerobic degradation of hydrocarbons and aromatic organic compounds^64–70^. Similarly, when we extended our search to a human metagenome-assembled genomes dataset that contains >150k genomes of human-associated microbes, all high homology hits (>90% amino acid identity) were identified in the genomes of *Enterocloster* species (**Figure S3E**). Altogether, this analysis suggests that the distribution of Ucdh is limited to a few *Enterocloster* species in the human gut.

### Metagenomic abundance of the *eam* gene cluster correlates with urolithin A levels and gut inflammation

Having characterized the biochemical activities of urolithin A-producing enzymes *in vitro*, we sought to evaluate their relevance *in vivo* by correlating fecal urolithin A levels with urolithin A-producing gene abundance in datasets from clinical studies. We chose to investigate a cross-sectional cohort of Crohn’s disease (CD), ulcerative colitis (UC), and non-inflammatory bowel disease (nonIBD) subjects from the Prospective Registry in IBD Study at MGH (PRISM)^71^. This cohort includes paired metagenomic and untargeted metabolomic data for stool samples from 155 subjects. We first quantified the metagenomic abundance of each urolithin A-producing gene (*eah*, *eadh1*, *eadh2*, and *ucdh*) separately. Since *eah*, *eadh1*, and *eadh2* are found in the same gene cluster, we averaged the abundance of these genes to calculate the Coriobacteriia <u>EA</u> <u>m</u>etabolism (*eam*) gene cluster abundance. Consistent with the high prevalence of the encoding species in the human gut^71^, the *eam* gene cluster and *ucdh* are present in the majority of stool metagenomes from healthy participants (98.2% for the *eam* gene cluster and 85.7% for *ucdh*). Next, we identified the mass peak corresponding to urolithin A in the untargeted metabolomics dataset by co-elution of stool metabolite samples with an authentic urolithin A standard (**Figure S4A** and **Figure S4B**). Upon multivariate linear regression analysis with age and medication use as co-variates, we found statistically significant positive correlations between the *eam* gene cluster abundance and urolithin A levels for the CD (adjusted *p* = 0.006) and UC subjects (adjusted *p* = 0.033) (**Figure 5A**). No significant correlation was found in the nonIBD subjects (adjusted *p* = 0.252), potentially due to its small sample size (n = 34). On the other hand, no significant correlations were observed between *ucdh* abundance and urolithin A levels in any cohort (**Figure 5A**). Though further studies are needed to understand this lack of correlation, our results revealed a possible connection between *in vivo* urolithin A levels and the Coriobacteriia *eam* gene cluster.

**Figure 5.**
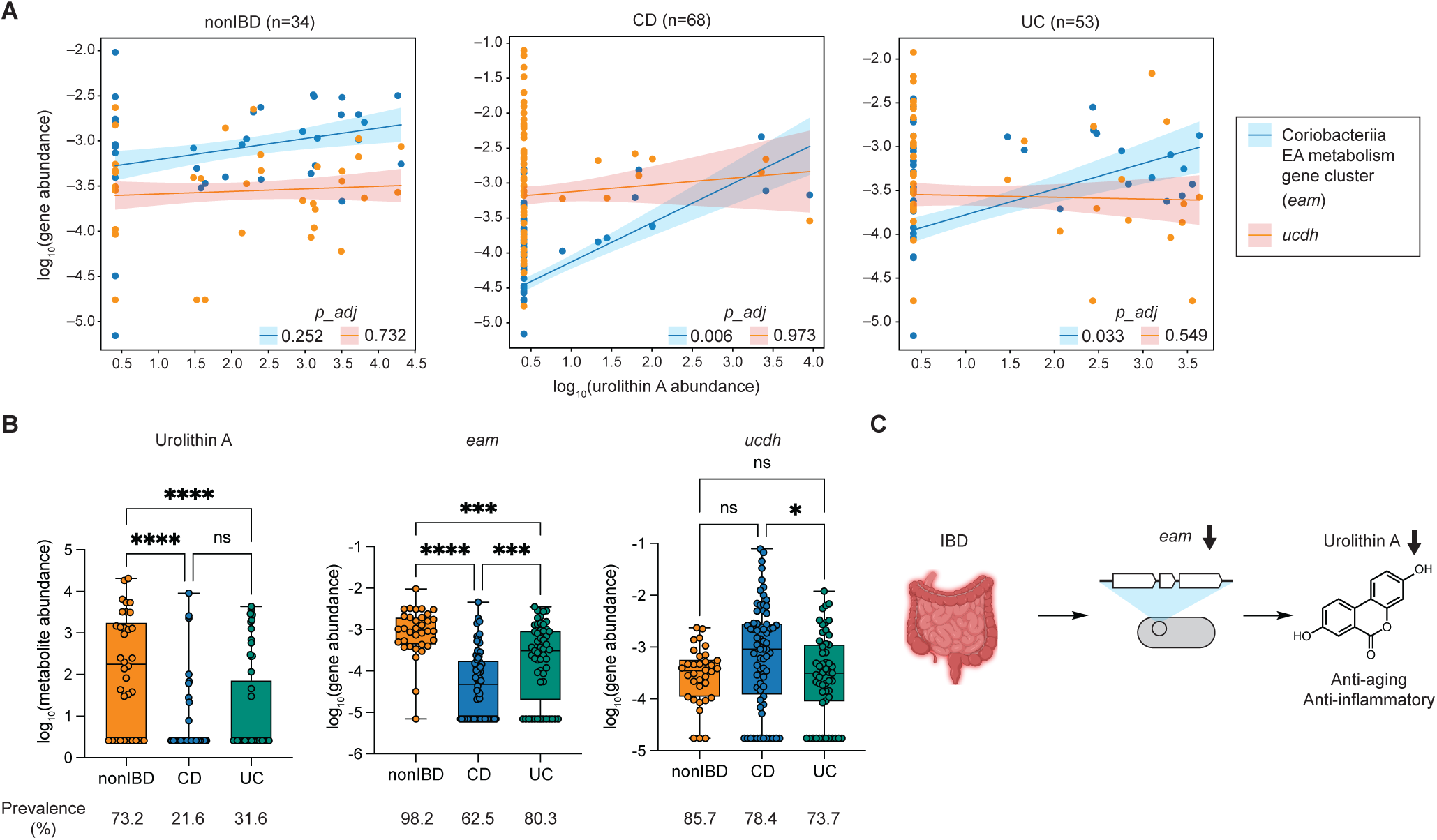
Metagenomic abundance of the *eam* gene cluster correlates with urolithin A levels and gut inflammation. **(A)** Scatter plots showing multivariate linear regression analyses between urolithin A metabolomic abundance and the *eam* gene cluster (average of *eah*, *eadh1*, and *eadh2*) or *ucdh* metagenomic abundances. Lines represent trend, and colored areas represent standard error. Adjusted *p* values were calculated using Benjamini-Hochberg correction with a target FDR of 0.05. **(B)** Boxplots of urolithin A concentration, Coriobacteriia gene cluster abundance, and *ucdh* abundance across host phenotypes. One-way ANOVA, Tukey’s multiple comparison. ns, non-significant; *, 0.01 < *p* value < 0.05; ***, 0.0001 < *p* value < 0.001; ****, *p* value <0.0001 **(C)** Schematic of a potential mechanistic link connecting IBD, the Coriobacteriia gene cluster, and urolithin A. IBD, inflammatory bowel disease; CD, Crohn’s disease; UC, ulcerative colitis.

We then compared metabolite and gene abundances between host phenotypes considering inflammation alters the production of gut bacterial metabolites^71–74^. Specifically, we postulated that reactive oxygen species generated during gut inflammation may negatively affect EA-metabolizing anaerobic bacteria and enzymes^75,76^. We found that urolithin A levels and the *eam* gene cluster abundance were lower in IBD compared to nonIBD controls (**Figure 5B**). Additionally, whereas urolithin A was present in 73.2% of nonIBD controls, only 21.6% of CD and 31.6% of UC samples had detectable levels of the metabolite. Furthermore, the *eam* gene cluster was detected more frequently in stool metagenomes from nonIBD subjects (98.2%) compared to CD (62.5%) and UC (80.3%). On the other hand, *ucdh* abundance did not show a significant decrease in IBD subjects. Since urolithin A has anti-inflammatory and anti-aging effects, this reduction in levels of urolithin A and producing enzymes may suggest IBD patients experience reduced health benefits from EA consumption (**Figure 5C**). Further investigation is required to test the causality of this suggested mechanistic connection.

## Discussion

In this study, we identified gut bacterial enzymes (a decarboxylase [Eah], three catechol dehydroxylases [Eadh1/2/3], and a xanthine oxidase enzyme [Ucdh]) that produce the anti-inflammatory urolithin A and other urolithins from EA. The conversion of EA to urolithin A represents the first example of the two classes of phenol dehydroxylases functioning in tandem (**Figure 6A**), expanding our fundamental understanding of gut bacterial enzymes involved in phenol dehydroxylation, a key transformation in the anaerobic metabolism of aromatic compounds. Considering their mutually exclusive regioselectivity (C4, C8, C10-OHs removed by catechol dehydroxylases and C9-OH removed by Ucdh) and differing substrate requirements, we suspect that these two enzyme classes employ distinct mechanisms to accomplish seemingly similar reactions. The mechanism of Ucdh may be analogous to that of 4-HBCR, for which two potential mechanisms have been proposed: a Birch-type one-electron process^57,58^ and a two-electron process^77^ (**Figure 6B** and **Figure S5A**). For both mechanisms, the electron-withdrawing thioester group of 4-hydroxybenzoyl-CoA is suggested to play an essential role in stabilizing the anion or radical intermediate generated during catalysis. Notably, the C9-OH of the urolithins is the only hydroxyl group on this scaffold positioned *para* to an electron-withdrawing functional group, perhaps explaining why the other hydroxyl groups are not removed by enzymes from this class (**Figure 6B**). Moreover, these proposed mechanisms do not require the departing hydroxyl group to be part of a catechol, consistent with our findings. In contrast, the proposed mechanism for catechol dehydroxylases requires an adjacent hydroxyl group, which undergoes tautomerization to prime the substrate for dehydroxylation (**Figure S5B**)^44,78^. Further studies are needed to understand the mechanistic basis for regioselectivities of the two molybdenum-dependent dehydroxylase classes. Recently, C3-OH dehydroxylated urolithins were identified in human stool samples, suggesting the presence of as-yet-uncharacterized phenol dehydroxylase targeting the C3-OH of urotlihins^79^. Our findings suggest the responsible enzyme is likely an uncharacterized catechol dehydroxylase because C3-OH is not positioned *para* to an electron-withdrawing group.

**Figure 6.**
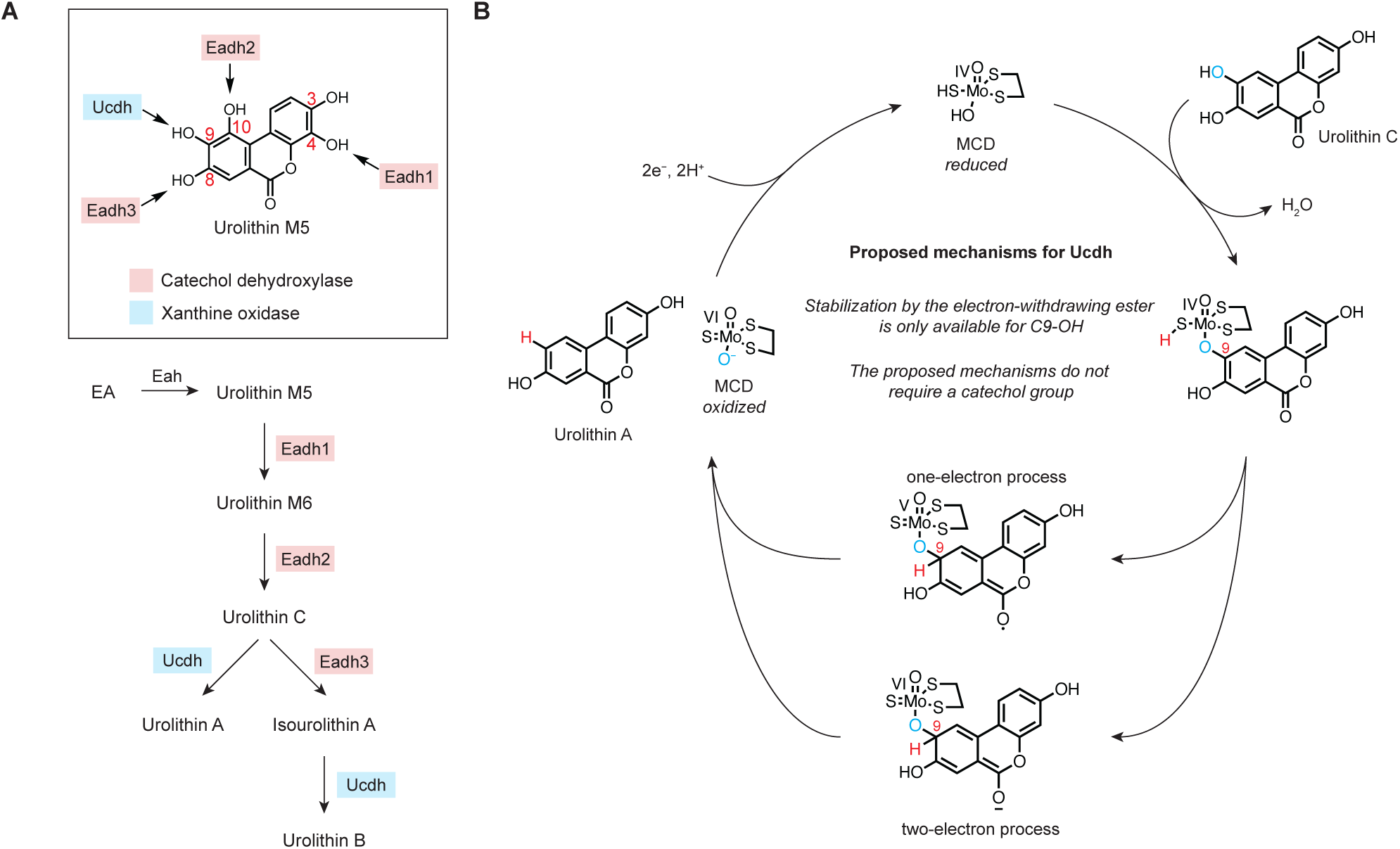
Proposed mechanisms for Ucdh may explain regioselectivity of urolithin dehydroxylation. **(A)** Mutually exclusive regioselectivity of Ucdh and Eadh1/2/3 (top) and the assignment of the complete EA metabolic pathways to urolithin A and urolithin B. **(B)** The lactone functional group of the urolithins may stabilize proposed intermediates during Ucdh catalysis.

Whereas other catechol dehydroxylases process *para* hydroxyl groups of mono-alkyl catechol substrates (**Figure 1C**), Eadh1/2/3 remove chemically distinct hydroxyl groups, including the sterically hindered C4-OH and C10-OH of urolithins, expanding the substrate scope of this enzyme class. The removal of such congested hydroxyl groups by this enzyme class is intriguing, especially given that catechol dehydroxylase substrates have been proposed to bind within a narrow substrate funnel^78^. Additionally, our kinetic experiments showed a strong substrate preference for each catechol dehydroxylase, explaining the sequence of the metabolic pathway. For example, the >60-fold higher catalytic efficiency of Eadh1 over Eadh2 toward urolithin M5 aligns with the predominant production of urolithin M6 over urolithin D in culture^30,32^. Similarly, previous reports analyzing urolithins in human stool samples did not detect urolithin D^79,80^. Interestingly, urolithin D was observed as a major intermediate in Iberian pigs, suggesting that enzymes encoded in pig microbiomes may have different substrate preferences^56^. In summary, our work expands the substrate scope of the catechol dehydroxylase class and connects enzyme kinetic parameters with the observed metabolic pathway of EA in humans.

Our transcriptomic and biochemical experiments linked the critical transformation in urolithin A formation, dehydroxylation of urolithin C C9-OH by *E. bolteae*, to the xanthine oxidase family enzyme Ucdh. We note that a recent preprint also linked this enzyme to urolithin C dehydroxylation using substrate-induced expression^81^. However, their attempts to support this functional assignment were limited by the minimal activity and poor solubility of heterologously expressed Ucdh. Here, we addressed this challenge by using the TP1000 *E. coli* strain, which overproduces the MCD cofactor required by xanthine oxidase enzymes. Co-expression of Ucdh with its putative chaperone XdhC significantly improved activity in TP1000 cell suspensions, a strategy that was previously shown to increase activity of a xanthine dehydrogenase^59^. After 4-HBCR, Ucdh is the second enzyme found to perform reductive dehydroxylation within the xanthine oxidase family, whose members catalyze mostly oxidative reactions^77^. Considering the phylogenetic relatedness of Ucdh and 4-HBCR, we hypothesize that these two enzymes share traits that allow this chemical reversal from oxidation to reduction. Although the molecular basis for this shift in reactivity remains unclear, the discovery of Ucdh implies that additional reductive enzymes remain to be uncovered within the xanthine oxidases. For example, a recent study found that human gut species encoding an uncharacterized xanthine oxidase enzyme reduce uric acid to xanthine using the reverse reaction of xanthine dehydrogenase^82^. However, this enzyme’s function has not been biochemically characterized. Another recent study identified and characterized a gut bacterial xanthine oxidase homolog that reductively cleaves an aromatic C–S bond in the ergothioneine catabolic pathway^83^, further expanding the repertoire of reductive chemistry catalyzed by this enzyme family. Our results, including the anaerobic protein expression system for Ucdh, will enable future efforts to biochemically characterize Ucdh and additional reductive xanthine oxidases.

Leveraging our knowledge of the genes and enzymes involved in EA metabolism, we found correlations between *in vivo* urolithin A levels and the abundance of urolithin A-producing genes in human gut microbiomes. Interestingly, levels of urolithin A were positively correlated with the abundance of the urolithin C-producing Coriobacteriia (*eam*) gene cluster. This correlation may be related to a previously reported positive correlation between *Gordonibacter* species levels and urolithin A production in healthy volunteers, although we only observed the positive correlation in IBD patient samples^28^. In contrast, levels of urolithin A were not correlated with *ucdh*, which may arise from discrepancies between *in vivo* metabolic activity and metagenomic abundance. Alternatively, we cannot rule out the possibility that some individuals might not have consumed EA or the presence of convergently evolved gut bacterial enzymes that produce urolithin A. Some previous reports described the isolation of urolithin A-producing gut bacterial strains from the *Enterococcus*^84^, *Lactococcus*^85^, or *Bifidobacterium* genera^86^. Since the EA-metabolizing genes we discovered are restricted to Coriobacteriia and *Enterocloster*, strains from other genera may use different enzymes to metabolize EA. However, to our knowledge, these metabolic activities have not been reproduced, and these strains are not commercially available nor fully sequenced. Additional efforts are thus required to validate their roles in EA metabolism and to identify responsible bacterial enzymes. Finally, we observed a decrease in both urolithin A levels and *eam* gene cluster abundance in the IBD cohort compared to nonIBD controls, suggesting a potential route by which host inflammation alters gut microbiome composition and impacts metabolite production. Lowered urolithin A levels in subjects with IBD may indicate they derived reduced benefits from consuming EA-rich foods. However, we acknowledge that altered levels of urolithin A could also arise from other phenotypic differences, including differences in dietary habits. Mechanistic investigation linking IBD, EA-metabolizing gene levels, and urolithin A production requires further testing in model organisms or human clinical cohorts. These results demonstrate the utility of applying obtained knowledge of gut bacterial metabolic genes and enzymes to human multi-omics datasets.

## Acknowledgments

This work was funded by the National Science Foundation (Alan T. Waterman Award to E.P.B., CHE-20380529) and the Biocodex Microbiota Foundation. E.P.B. is a Howard Hughes Medical Institute Investigator. M.B. acknowledges support from the Kwanjeong Educational Foundation. We thank Harvard Research Computing for computational resources, maintenance, and support. We thank Dr. Richard Y. Liu, Dr. Francisco A. Tomás-Barberán and Dr. Eric Franzosa for insightful discussions. We thank Dr. Gil Namkoong and Rohan Narayan for critical reading of the manuscript. This article is subject to HHMI’s Open Access to Publications policy. HHMI lab heads have previously granted a nonexclusive CC BY 4.0 license to the public and a sublicensable license to HHMI in their research articles. Pursuant to those licenses, the author-accepted manuscript of this article can be made freely available under a CC BY 4.0 license immediately upon publication.

## Author contributions

M.B., V.M.R., and E.P.B. conceptualized the project. M.B., X.D., J.A.P., C.C., F.I., and V.M.R. performed experiments and analyzed the data. M.B. and E.P.B. wrote the manuscript. All authors provided critical feedback on the manuscript.

## Data availability

Raw RNA-seq data of *Gs* 28C and *E. bolteae* have been deposited in the Sequence Read Archive under BioProject accession number: PRJNA1110272.

## Materials and Methods

### General materials and methods

The following chemicals were used in this study: ellagic acid (EA) (Sigma-Aldrich, catalog# E2250-1G), urolithin M5 (Toronto Research Chemicals Inc., catalog# TRC-U847035-1mg), urolithin M6 (Toronto Research Chemicals Inc., catalog# TRC-U847040-10mg), urolithin D (Toronto Research Chemicals Inc., catalog# TRC-U847020-5mg), urolithin E (Toronto Research Chemicals Inc., catalog# TRC-U847030-10mg), urolithin M7 (Toronto Research Chemicals Inc., catalog# TRC-U847045-2.5mg), urolithin C (Sigma-Aldrich, catalog# SML3047), urolithin A (AstaTech, catalog# Y10574), isourolithin A (AstaTech, catalog# F51950), urolithin B (Sigma-Aldrich, catalog# OTV000001), L-arginine monohydrochloride (Sigma-Aldrich, catalog# A5131-500G), L-cysteine hydrochloride monohydrate (Sigma-Aldrich, catalog# C7880-100G), sodium formate (Sigma-Aldrich, catalog# 247596-100G), methyl viologen (MV) (Sigma-Aldrich, catalog# 856177-1g), sodium dithionite (NaDT) (Sigma-Aldrich, catalog# 157953-5G), sodium molybdate dihydrate (Sigma-Aldrich, catalog# 243655-100G), dimethylformate (DMF) (Sigma-Aldrich, catalog# 319937-4L), Isopropyl β-D-1-thiogalactopyranoside (IPTG, ultra-pure grade) (Teknova, catalog# I3325), 4-Isopropylbenzoic acid (cumate) (Ambeed, catalog# A255165), kanamycin sulfate (VWR, catalog# 75856-686), ampicillin (IBI scientific, catalog# IB02040), sodium nitrate (Sigma-Aldrich, catalog# S5506), ammonium ferric citrate (Sigma-Aldrich, catalog# F5879), sodium fumarate (Sigma-Aldrich, catalog# F1506), SIGMAFAST protease inhibitor tablets (Sigma-Aldrich, catalog# S8830), DNase (Sigma-Aldrich, catalog# DN25-1G), lysozyme (Sigma-Aldrich, catalog# L6876-5G), β-Nicotinamide adenine dinucleotide reduced form disodium salt trihydrate (NADH) (Research Products International, catalog# N20100-1.0). LC–MS grade acetonitrile and methanol for LC–MS analyses were purchased from Honeywell Burdick and Jackson. LC–MS grade formic acid was purchased from Sigma-Aldrich (catalog# 5330020050). Brain–Heart Infusion (BHI) broth was purchased from Becton Dickinson (catalog# 211060). Luria-Bertani (LB) medium was purchase from Research Products International (catalog# L24060). TRIzol was purchased from Invitrogen (catalog# 15596026).

10-beta competent *E. coli* (New England Biolabs, C3019H) was routinely used for cloning and DNA construction. *Gordonibacter sp.* 28C, *Paraeggerthella hongkongensis* RC2/2 A, and *Eggerthella* strains were obtained from the Peter Turnbaugh lab at the University of California, San Francisco. *Adlercreutzia equolifaciens* DSM 19450, *Senegalimassilia anaerobia* DSM 25959, *Gordonibacter urolithinfaciens* DSM 27213, *Gordonibacter pamelaeae* 7-10-1b, *Ellagibacter isourolithinifaciens* DSM 104140, *Rubneribacter badeniensis* DSM 105129, and *Raoultibacter massiliensis* DSM 103407 were purchased from Leibniz Institute DSMZ. *E. coli* TP1000 strain was obtained as a generous gift from Prof. Kurt Warnhoff at Sanford Research.

To prepare electrocompetent *G. urolithinfaciens* cells for plasmid transformation, a turbid 48-hour starter culture of *G. urolithinfaciens* strain in BHI medium was inoculated 1:50 into 40 mL of BHI medium supplemented with 1% L-arginine monohydrochloride (w/v%). When the culture reached optical density at 600 nm (OD600) of 0.3, the cells were pelleted by centrifugation, and the supernatant was removed. The cell pellet was washed with 10 mL of ice-cold sterile water three times and then with 5 mL of deoxygenated sterile 10% (v/v) glycerol aqueous solution. Finally, the cell pellet was resuspended in 2.5 mL of deoxygenated sterile 10% (v/v) glycerol solution and subdivided into 100 µL aliquots, which were flash frozen and stored at –80 °C until use.

To prepare electrocompetent *E. coli* BL21(DE3) or TP1000 cells for plasmid transformation, a turbid overnight culture of *E. coli* strains in LB medium was inoculated 1:100 into 100 mL of LB medium. When the culture reached OD600 of 0.4–0.6, the cells were pelleted by centrifugation at 4 °C, and the supernatant was removed. The cell pellet was washed with 40 mL of ice-cold sterile 10% (v/v) glycerol aqueous solution five times. Finally, the cell pellet was resuspended in 1 mL of sterile 10% (v/v) glycerol solution and subdivided into 50 µL aliquots, which were flash frozen and stored at –80 °C until use.

Unless otherwise noted, all bacterial culturing work was performed in an anaerobic chamber (Coy Laboratory Products) under an atmosphere of 2–4% hydrogen, 2–4% carbon dioxide, and nitrogen as the balance. Hungate tubes were used for anaerobic culture unless otherwise noted (Chemglass, catalog# CLS-4209–01). The human gut Coriobacteriia species were grown at 37 °C on BHI supplemented with 1% L-arginine monohydrochloride (w/v%), 0.05% L-cysteine monohydrochloride (w/v%), 10 mM sodium formate (BHIrcf medium) unless otherwise noted.

Bioinformatics analyses were done on the Harvard Faculty of Arts & Sciences Research Computing Cluster.

### Coriobacteriia screening for EA metabolizers

Turbid 48-hour starter cultures of Coriobacteriia species in BHI medium were diluted 1:100 into 200 µL of BHIrcf medium supplemented with 100 µM EA in triplicates on a 96-well plate. The plate was then sealed, and the cultures were anaerobically grown at 37 °C for 72 hours. For LC–MS/MS analysis of dehydroxylation activity, 40 µL of each supernatant was diluted into 160 µL of LC–MS grade methanol, and the precipitates were removed by centrifugation before injection onto LC–MS/MS, as described below.

### Substrate-dependent inducibility experiment

#### Gordonibacter species 28C (cell lysate)

A turbid 48-hour starter culture of *Gs* 28C in BHI medium was diluted 1:100 into 5 mL of BHIrcf medium in six replicates. The cultures were anaerobically incubated at 37 °C. When OD600 of the cultures reached 0.2, 50 µL of a 10 mM solution of EA in DMF was added to three replicates (final EA concentration 100 µM), and 50 µL of DMF vehicle was added to the other three replicates. The cultures were further anaerobically incubated at 37 °C until OD600 reached 0.5, at which cells were pelleted by centrifugation. In an anaerobic chamber, the cell pellets were washed with 5 mL of 1X phosphate-buffered saline and re-suspended in 1 mL of lysis buffer (20 mM Tris, pH 7.5, 500 mM NaCl, 10 mM MgSO4, 1 mM CaCl2, 0.1 mg/mL DNase, 0.5 mg/mL lysozyme, 1 tablet/50 mL SIGMAFAST protease inhibitor). Cells were lysed via sonication (Branson Sonifier 450, 25% amplitude, 10 seconds on, 40 seconds off, 2 min total sonication time). The lysates were then clarified via centrifugation (20k RCF, 15 mins) and were subjected to the activity assay. In a 96-well plate, 1 µL of a 10 mM solution of EA in DMF (100 µM final concentration) or vehicle and 2 µL of 50 mM MV and 50 mM NaDT (1 mM final concentration) were added to 95 µL of cell lysate in triplicates. The plate was then sealed and left in the anaerobic chamber at room temperature for 48 hours. 40 µL of each reaction mixture was diluted into 160 µL of LC–MS grade methanol, and the precipitates were removed by centrifugation before injection onto LC–MS/MS, as described below.

#### Enterocloster bolteae DSM 15670 (cell suspension)

A turbid 48-hour starter culture of *E. bolteae* DSM 15670 in BHI medium was diluted 1:100 into 10 mL of BHI medium in six replicates. The cultures were anaerobically incubated at 37 °C. When OD600 of the cultures reached 0.3, 100 µL of a 10 mM solution of urolithin C in DMF was added to three replicates (100 µM final concentration), and 50 µL of vehicle was added to the other three replicates. The cultures were further anaerobically incubated at 37 °C until OD600 reached 0.9, at which cells were pelleted by centrifugation. In an anaerobic chamber, the cell pellets were washed with 5 mL of 1X phosphate-buffered saline twice and re-suspended in 1 mL of 1X phosphate-buffered saline. In a 96-well plate, 1 µL of a 10 mM solution of urolithin C in DMF (100 µM final concentration) or vehicle was added to 99 µL of cell suspension in triplicates. The plate was then sealed and left in the anaerobic chamber at room temperature for 6 hours. 40 µL of each reaction mixture was diluted into 160 µL of LC–MS grade methanol, and the precipitates were removed by centrifugation before injection onto LC–MS/MS, as described below.

#### RNA sequencing

(*Gs* 28C) A turbid 48-hour starter culture of *Gs* 28C in BHI medium was diluted 1:100 into 40 mL of BHIrcf medium in six replicates. The cultures were anaerobically incubated at 37 °C. When the OD600 of the cultures reached 0.2, 400 µL of a 10 mM solution of EA in DMF (100 µM final concentration) or vehicle was added to the cultures in three replicates. The cultures were further anaerobically incubated at 37 °C until OD600 reached 0.5, at which cells were pelleted by centrifugation and resuspended in 500 µL of Trizol. Total RNA was isolated by first bead beating to lyse cells and then using the Zymo Research Direct-Zol RNA MiniPrep Plus kit (Catalog # R2070) according to the manufacturer’s protocol. Illumina cDNA libraries were generated using a modified version of the RNAtag-Seq protocol^87^. Briefly, 500 ng of total RNA was fragmented, depleted of genomic DNA, and dephosphorylated prior to its ligation to DNA adapters carrying 5’-AN8-3’ barcodes with a 5’ phosphate and a 3’ blocking group. Barcoded RNAs were pooled and depleted of rRNA using the RiboZero rRNA depletion kit (Epicentre). These pools of barcoded RNAs were converted to Illumina cDNA libraries in three main steps: (i) reverse transcription of the RNA using a primer designed to the constant region of the barcoded adaptor; (ii) addition of a second adapter on the 3’ end of the cDNA during reverse transcription using SmartScribe RT (Clonetech) as described^87^; (iii) PCR amplification using primers that target the constant regions of the 3’ and 5’ ligated adaptors and contain the full sequence of the Illumina sequencing adaptors. cDNA libraries were sequenced on Illumina HiSeq 2500. For the analysis of RNAtag-Seq data, reads from each sample in the pool were identified based on their associated barcode using custom scripts, and up to one mismatch in the barcode was allowed with the caveat that it did not enable assignment to more than one barcode. Barcode sequences were removed from the first read as were terminal Gs from the second read that may have been added by SMARTScribe during template switching.

(*Eb* DSM 15670) A turbid 48-hour starter culture of *Gs* 28C in BHI medium was diluted 1:100 into 10 mL of BHIrcf medium in six replicates. The cultures were anaerobically incubated at 37 °C. When the OD600 of the cultures reached 0.5, 100 µL of a 10 mM solution of EA in DMF (100 µM final concentration) or vehicle was added to the cultures in three replicates. The cultures were further anaerobically incubated at 37 °C until OD600 reached 0.9, at which cells were pelleted by centrifugation and resuspended in 500 µL of Trizol. Total RNA was extracted using Trizol following manufacturer’s instructions (ThermoFisher Scientific, Waltham, MA, USA). RNA samples were quantified using Qubit 2.0 Fluorometer (ThermoFisher Scientific, Waltham, MA, USA) and RNA integrity was checked with 4200 TapeStation (Agilent Technologies, Palo Alto, CA, USA). rRNA depletion sequencing library was prepared by using three probes from QIAGEN FastSelect rRNA 5S/16S/23S Kit (Qiagen, Hilden, Germany), respectively. RNA sequencing library preparation uses NEBNext Ultra II RNA Library Preparation Kit for Illumina by following the manufacturer’s recommendations (NEB, Ipswich, MA, USA). Briefly, enriched RNAs are fragmented for 15 minutes at 94 °C. First strand and second strand cDNA are subsequently synthesized. cDNA fragments are end repaired and adenylated at 3’ends, and universal adapters are ligated to cDNA fragments, followed by index addition and library enrichment with limited cycle PCR. Sequencing libraries were validated using the Agilent Tapestation 4200 (Agilent Technologies, Palo Alto, CA, USA), and quantified using Qubit 2.0 Fluorometer (ThermoFisher Scientific, Waltham, MA, USA) as well as by quantitative PCR (KAPA Biosystems, Wilmington, MA, USA). The sequencing libraries were multiplexed and clustered onto 1 lane of a flowcell. After clustering, the flowcell was loaded onto the Illumina HiSeq instrument (4000 or equivalent) according to manufacturer’s instructions. The samples were sequenced using a 2x150bp Paired End (PE) configuration. Image analysis and base calling were conducted by the Illumina Control Software. Raw sequence data (.bcl files) generated was converted into fastq files and de-multiplexed using Illumina bcl2fastq 2.17 software. One mis-match was allowed for index sequence identification.

#### Differential expression analysis

Kneaddata v0.10.0 (--bypass-trf) was used to clean up raw sequencing data by removing short or poor-quality reads and clipping off Illumina sequencing adaptor sequences. Bowtie2 v2.5.1^88^ and HTSeq 2.0 v0.11.2^89^ were used to align reads to the genome of the sequenced organisms and to count the number of mapped reads onto each protein-encoding gene. DESeq2 v1.44.0^90^ was used to compute fold change, mean of normalized counts, and adjusted p-value of each protein-encoding gene. Variance among genes with low expression levels was stabilized by applying lfcShrink (coef=2, type=“apeglm”)^91^. The plotMA function was used to visualize the results^92^.

#### Plasmid construction

Plasmid construction was carried out using standard molecular biology techniques. (Eah) To construct a plasmid for Eah expression, we first amplified the Eah gene from *Gs* 28C gDNA using primers oMB-Eah-F and oMB-Eah-R (with Gibson Assembly overhangs) and the vector backbone from a pET28a vector using primers oMB-pET28a-F and pMB-oET28a-R. The vector backbone was then gel purified and DpnI digested. The amplified gene fragment and the vector backbone were ligated using Gibson Assembly (pMB-pET28a-Eah). The primers were designed to introduce an N-terminal His6 tag into Eah. (Eadh1, Eadh2, Eadh3) To construct a cumate-inducible expression vector for catechol dehydroxylases in *G. urolithinfaciens*, we first linearized a previously prepared vector pXD80GHcdh3-pXD68Kan2-Pct3 using a pair of restriction enzymes NsiI and SpeI. The *G. urolithinfaciens* genomic regions containing protein-encoding genes for Eadh1 and Eadh2 subunits, with an insertion of a sequence encoding N-terminal His6 tag to subunit A, were synthesized by Genewiz. Likewise, the *E. isourolithinifaciens* genomic region containing Eadh3 subunits, with an insertion of a sequence encoding N-terminal His6 tag into subunit A, was synthesized by Genewiz. Gibson assembly overhang sequences were introduced by amplifying the synthesized gene sequences using primer pairs oMB-Eadh1-F/R, oMB-Eadh2-F/R, oMB-Eadh3-F/R for Eadh1, Eadh2, and Eadh3, respectively. The amplified gene fragments were inserted into the linearized pXD68Kan2 backbone using Gibson Assembly (pMB-pXD80-Eadh1, pMB-pXD80-Eadh2, pMB-pXD80-Eadh3). (Ucdh) To construct a plasmid for Ucdh expression, we first linearized pTrcHis2A using a pair of restriction enzymes BamHI and HindIII. Ucdh subunits were amplified from *E. bolteae* gDNA as two gene fragments using primer pairs oMB-Ucdh-F/oMB-Ucdh-His-R and oMB-Ucdh-His-F/oMB-Ucdh-noXdhC-R. The primers were designed to insert a sequence encoding N-terminal His6 tag into subunit C The two amplified gene fragments and the linearized pTrcHis2A backbone were ligated using Gibson Assembly (pMB-pTrcHis2A-Ucdh). To construct a plasmid for Ucdh and XdhC co-expression, Ucdh subunits were amplified from pMB-pTrcHis2A-Ucdh using a primer pair oMB-Ucdh-F/R, and XdhC was amplified from *E. bolteae* gDNA using a primer pair oMB-XdhC-F/R. The amplified gene fragments and the linearized pTrcHis2A backbone were ligated using Gibson Assembly (pMB-pTrcHis2A-Ucdh-XdhC).

#### Expression of Eah in E. coli BL21(DE3)

The chemically competent *E. coli* BL21(DE3) was transformed with 100 ng of pMB-pET28a-Eah, and transformants were selected on a LB agar plate supplemented with kanamycin (50 µg/mL). A single colony was picked and inoculated into LB medium supplemented with kanamycin (50 µg/mL) and grown for overnight at 37 °C. The saturated culture was inoculated 1:200 into 1 L of fresh LB medium supplemented with kanamycin (50 µg/mL) in a 2.8 L Erlenmeyer baffled flask and grown at 37 °C at 180 rpm for 3 hours, at which OD600 reached 0.5. IPTG was added to the final concentration of 0.25 mM. The culture was then grown at 18 °C at 180 rpm for 16 hours. Cell pellets were harvested by centrifugation at 6,000 × g for 20 min at 4 °C and used for protein purification immediately.

#### Expression of catechol dehydroxylases in G. urolithinfaciens

For transformation, electrocompetent *G. urolithinfaciens* cells were electroporated using a MicroPulser Electroporator (BioRad) with 100–1,000 ng of pMB-pXD80-Eadh1/2/3 at 2.5 kV voltage in 1-mm gap width electroporation cuvettes (VWR). 1 mL of pre-reduced BHI medium supplemented with 1% L-arginine monohydrochloride (w/v%) and 10 mM sodium formate was immediately added to the electroporated cells and transferred to 1.7 mL Eppendorf tubes. These tubes were brought to anaerobic chamber (Coy Laboratory Products) and incubated at 37 °C for 3 hours. The cultures were then plated onto BHI agar plates supplemented with 1% L-arginine monohydrochloride (w/v%), 10 mM sodium formate, and kanamycin (100 µg/mL) and grown anaerobically for 3–5 days at 37 °C. A colony was picked and inoculated into BHIrcf medium with kanamycin (100 µg/mL) and grown for 2 days at 37 °C anaerobically. The saturated culture was inoculated 1:25 into 2 L of fresh BHIrcf medium with kanamycin (100 µg/mL) and cumate (100 µM) in a 2.8 L anaerobic Erlenmeyer baffled flask. The culture was grown at 37 °C anaerobically for 20–24 hours without shaking. Cell pellets were harvested by centrifugation at 6,000 × g for 20 min at 4 °C and used for protein purification immediately.

#### Expression of Ucdh in E. coli TP1000

Chemically competent *E. coli* TP1000 was transformed with 100 ng of pMB-pTrcHis2A-Ucdh-XdhC, and transformants were selected on a LB agar plate supplemented with kanamycin (100 µg/mL) and ampicillin (100 µg/mL). A single colony was picked and inoculated into LB medium supplemented with kanamycin (100 µg/mL) and ampicillin (100 µg/mL) and grown for overnight at 37 °C. The saturated culture was inoculated 1:200 into 2 L of fresh LB medium supplemented with 1 mM sodium molybdate, 2 mM ammonium ferric citrate, and ampicillin (100 µg/mL) in a 2.8 L anaerobic Erlenmeyer baffled flask and grown at 37 °C at 180 rpm for 3 hours, at which OD600 reached 0.6. The culture was then sparged with nitrogen gas for 30 min, after which 25 mM sodium fumarate and 10 mM sodium nitrate were added. After the additional sparging with nitrogen gas for 30 min, 25 µM IPTG was added, and the flask was tightly capped. The culture was then grown anaerobically at 18 °C for 16 hours without shaking. Cell pellets were harvested by centrifugation at 6,000 × g for 20 min at 4 °C and used for protein purification immediately.

#### Enzyme Purification

All subsequent steps were performed at 4 °C unless otherwise specified. Eah was purified aerobically whereas catechol dehydroxylases and Ucdh were purified anaerobically. Cell pellets were resuspended at 5 mL per g cell paste in lysis buffer (50 mM HEPES, 30 mM imidazole, 250 mM NaCl, 2 mg/mL lysozyme, pH 8). Cells were lysed by sonication using a ½ inch horn at 35% amplitude for 6 min (10 sec on followed by 40 sec off) while being kept in an ice water bath. Lysate was centrifuged at 14,000 × g for 50 min at 4 °C to separate soluble and insoluble fractions. 2 mL bed volume of Ni-NTA resin was equilibrated with 20 mL of lysis buffer and was then mixed with soluble lysate followed by 1 hour incubation with constant agitation. The lysate was then loaded onto a column by gravity flow. The column was washed with 10 column volumes of wash buffer (50 mM HEPES, 30 mM imidazole, 250 mM NaCl, pH 8). Protein was eluted from the column with elution buffer (50 mM HEPES, 250 mM imidazole, 250 mM NaCl, pH 8). Combined elution fractions containing protein were concentrated using an Ultra-15 Centrifugal Filters centrifugal concentrator with a 30 kDa MWCO membrane (Amicon) and desalted using a disposable PD-10 desalting column (cytiva). The desalted proteins were again concentrated using a new centrifugal concentrator with a 30 kDa MWCO membrane (Amicon). Protein concentrations were estimated using a NanoDrop 2000 UV-Vis Spectrophotometer (Thermo Scientific) using extinction coefficient at a wavelength of 280 nm (ε280) calculated by ExPASY ProtParam as follows: Eadh1A(NHis6)B (ε280 = 227,965 M^−1^ cm^−1^), Eadh2A(NHis6)B (ε280 = 215,725 M^−1^ cm^−1^), Eadh3A(NHis6)B (ε280 = 229,955 M^−1^ cm^−1^). Protein concentrations of Eah(NHis6) and UcdhABC(NHis6) were calculated using a Bradford assay. The protein samples were flash frozen and stored at –80 °C until use.

#### Activity assay of E. coli TP1000 expressing Ucdh in cell suspension

A single colony of TP1000 strains containing either of pTrcHis2A (empty vector), pMB-pTrcHis2A-Ucdh, or pMB-pTrcHis2A-Ucdh-XdhC was picked and inoculated into 5 mL LB medium with 100 µg/mL kanamycin and 100 µg/mL ampicillin and grown for overnight at 37 °C. The saturated cultures were inoculated 1:100 into 10 mL of fresh LB medium supplemented with 1 mM sodium molybdate, 2 mM ammonium ferric citrate, 10 mM sodium nitrate, 25 µM IPTG, and 100 µg/mL ampicillin in 14 mL culture tubes in triplicates. The cultures were brought into an anaerobic chamber and were grown at 37 °C for 24 hours under anaerobic conditions. In an anaerobic chamber, the cell pellets were washed with 1 mL of pre-reduced 1X phosphate-buffered saline twice and re-suspended into 1 mL of the 1X phosphate-buffered saline. In a 96-well plate, 1 µL of a 10 mM solution of urolithin in DMF (100 µM final concentration) or vehicle was added to 99 µL of cell suspension in triplicates. The plate was then sealed and left in the anaerobic chamber at rt for 24 hours. 40 µL of each reaction mixture was diluted into 160 µL of LC–MS grade methanol, and the precipitates were removed by centrifugation before injection onto LC–MS/MS, as described below.

### Phylogenetic analysis

#### Molybdenum enzymes from Coriobacteriia strains

Extracted amino acid sequences of proteins annotated as the catalytic subunit of molybdopterin-dependent oxidoreductases from the genomes of Coriobacteriia strains (*Gs* 28C, *G. pamelaeae* 3C, *G. pamelaeae* 7-10-1b, *G. urolithinfaciens* DSM 27213, and *E. isourolithinifaciens* DSM 104140) (total 305 sequences). These sequences were first grouped by 80% amino acid identity into 144 unique sequences using CD-HIT^93^. The unique sequences were then aligned using MAFFT-linsi v7.505^94^ and trimmed using trimal (v1.4.1, -gappyout)^95^. A maximum-likelihood phylogenetic tree was generated using iqtree2 (v2.1.3, 1000 ultrafast bootstraps)^96^. The tree was visualized using a free web-based tool, iTOL v6.5.8.

#### Xanthine oxidase

Protein sequences of the catalytic subunit from 61 biochemically characterized xanthine oxidase family enzymes (IPR016208) were downloaded from the Swiss-Prot database (accessed 2023/02/14). The protein sequences and Ucdh were then aligned using MAFFT-linsi v7.505^94^ and trimmed using tirmal (v1.4.1, -gappyout)^95^. A maximum-likelihood phylogenetic tree was generated using iqtree2 (v2.1.3, 1000 ultrafast bootstraps)^96^. The eukaryotic xanthine oxidases were served as outgroup for a tree of bacterial xanthine oxidases and Ucdh. The tree was visualized using a free web-based tool, iTOL v6.5.8.

### Metagenome-assembled genome search

#### Gordonibacter genes (Eadh1 and Eadh2)

The amino acid sequences of Eah and the catalytic subunits of Eadh1 and Eadh2 were used as queries for tBLASTn searches (search translated nucleotide databases using a protein query) with an e-value cutoff of <0.0001. The following six MAG datasets sampling microbial sequencing data from diverse environments ranging from mammalian-associated microbes^97–101^ to soils and oceans^102^ were searched. Hits with >80% amino acid identity to each query were considered homologs. The cutoff was chosen because sequences with lower homology may have different substrates^47,48^. Contigs containing the homologs were annotated using Kbase Prokka^103^.

#### Ucdh

The amino acid sequence of the catalytic subunit of Ucdh was used as a query for tBLASTn search with an e-value cutoff of <0.0001. The MAG dataset containing >150k MAGs assembled from human-associated microbial sequencing datasets was searched^97^.

#### Enzyme activity assays

All assays were performed in an anaerobic chamber containing N2 and <0.1 ppm O2 (MBRAUN). All assay concentrations are final concentrations.

#### Eah activity assay

1 µM Eah or blank enzyme storage buffer was added in triplicate to assay buffer containing 100 µM EA (50 mM HEPES, 250 mM NaCl, pH 7.0). The final reaction solution was sealed and left at rt, without shaking, for 20 hours. For LC–MS/MS analysis, 40 µL of the solution was first diluted 1:5 into LC–MS grade methanol, followed by centrifugation to remove precipitates before injection onto LC–MS/MS, as described below.

#### Catechol dehydroxylase activity assay

100 nM of each catechol dehydroxylase or blank enzyme storage buffer was added in triplicate to assay buffer containing 200 µM methyl viologen, 100 µM sodium dithionite, and 100 µM substrate (20 mM MOPS, 300 mM NaCl, pH 7.0). The final reaction solution was sealed and left at rt, without shaking, for 20–24 hours. For LC–MS/MS analysis, 40 µL of the solution was first diluted 1:5 into LC–MS grade methanol, followed by centrifugation to remove precipitates before injection onto LC–MS/MS, as described below.

#### Ucdh activity assay

2 µM Ucdh or blank enzyme storage buffer was added in triplicate to assay buffer containing 1 mM NADH and 100 µM substrate (20 mM MOPS, 300 mM NaCl, pH 7.0). The final reaction solution was sealed and left at rt, without shaking, for 20–24 hours. For LC–MS/MS analysis, 40 µL of the solution was first diluted 1:5 into LC–MS grade methanol, followed by centrifugation to remove precipitates before injection onto LC–MS/MS, as described below.

#### Catechol dehydroxylase kinetics

We adapted a continuous absorbance-based enzyme kinetics assay previously developed for catechol dehydroxylases^47^. All assay concentrations are final concentrations. Anoxic catechol dehydroxylases (Eadh1, Eadh2, or Eadh3) and solid substrates were brought into an anaerobic chamber containing N2 and <0.1 ppm O2 (MBRAUN). Substrates were dissolved in pre-reduced DMF. The anoxic buffer (50 mM MOPS, pH 7.0, 300 mM NaCl) contained 200 µM MV, 100 µM NaDT, 2-fold serial dilutions of urolithins (3.9 to 500 µM), and an enzyme. Enzyme concentrations were varied based on enzyme–substrate pairs: 10 nM for Eadh1–urolithin M5, 20 nM for Eadh1–urolithin E, and 200 nM for the other pairs. The initial rate over the first minute was measured using a continuous spectrophotometric assay. In this assay, the reductive dehydroxylation was coupled with the oxidation of MV^+×^ (blue, absorption maxima at 605 nm wavelength) to MV^2+^ (colorless). Each assay was performed using two independently prepared protein samples, with each sample tested in biological triplicates. Kinetic parameters were calculated using GraphPad Prism 10 (GraphPad Software). Cooperative kinetics were fit with the model v = (*k*cat ∗ [E]0 ∗ [S]) / (Km + [S]).

### LC–MS/MS methods

#### Method A (all metabolites except urolithin B)

Samples were analyzed using a Waters UPLC-QQQ Mass Spectrometer equipped with a CORTECS T3 column (Waters Corp.). The following chromatography conditions were used: Column temperature, 40 °C; Mobile phase A, water (0.1% formic acid); Mobile phase B, acetonitrile (0.1% formic acid); Gradient (percentage denotes the ratio of mobile phase A): 100%, 1 min; 100% to 40%, 1.5 min; 40% to 10%, 0.2 min; 10%, 0.3 min, 10% to 100%, 0.05 min; 100%, 0.35 min; Flowrate, 0.5 mL/min; Injection volume: 1.0 µL.

#### Method B (urolithin B)

Samples were analyzed using a Waters UPLC-QQQ Mass Spectrometer equipped with a ACQUITY UPLC BEH amide column (Waters Corp.). The following chromatography conditions were used: Column temperature, 40 °C; Mobile phase A, water (0.1% formic acid); Mobile phase B, acetonitrile (0.1% formic acid); Gradient (percentage denotes the ratio of mobile phase A): 10%, 0.5 min; 10% to 100%, 1.5 min; 100%, 0.5 min; 100% to 10%, 0.25 min; 10%, 0.65 min; Flowrate, 0.5 mL/min; Injection volume: 1.0 µL.

For MS/MS, the precursor and daughter ions were detected via Electrospray Ionization (ESI) in negative mode. Isomers with same MS/MS fragments were distinguished by comparing their retention times to the synthetic standards. The transitions and collision energy used to detect specific compounds of interest are listed. (1) EA: precursor ion *m/z*, 300.85; daughter ion *m/z*, 228.85; collision energy, 28 V; cone voltage, 92 V. (2) Urolithin M5: precursor ion *m/z*, 275.00; daughter ion *m/z*, 257.17; collision energy, 24 V; cone voltage, 82 V. (3) Urolithin M6, urolithin E and urolithin D: precursor ion *m/z*, 258.94; daughter ion *m/z*, 212.95; collision energy, 26 V; cone voltage, 92 V. (4) Urolithin C and urolithin M7: precursor ion *m/z*, 243.01; daughter ion *m/z*, 187.00; collision energy, 28 V; cone voltage, 30 V. (5) Urolithin A and isourolithin A: precursor ion *m/z*, 227.01; daughter ion *m/z*, 198.06; collision energy, 30 V; cone voltage, 52 V, (6) Urolithin B: precursor ion *m/z*, 211.21; daughter ion *m/z*, 167.09; collision energy, 24 V; cone voltage, 42 V

### Metagenomic analysis

The metagenomic dataset from the PRISM study was downloaded from the National Center for Biotechnology Information (PRJNA400072)^71^. Host-associated reads, low-quality reads, and adapter sequences were removed using KneadData v0.10.0 (default setting). Using a BLASTX DIAMOND search (e-value cutoff: 0.0001)^104^, reads were mapped on the following protein sequences (Eah [*Gs* 28C], Eadh1 [*Gs* 28C], Eadh2 [*Gs* 28C], and Ucdh [*Eb* DSM 15670]). To prevent non-specific read mapping, we also included UniRef50 sequences of the top 1000 BLAST hits of each query from UniProtKB (assessed Aug 2021) to the query list. Reads with >80% amino acid identity to each sequence were counted. If a read was mapped on multiple proteins, the read was counted as a hit for the protein that had the highest identity, which increases the specificity of the quantification. The number of hits was normalized first by the number of the total reads of each sample and the lengths of the proteins to read per kilobase million (RPKM). For further statistical analyses across samples, RPKM was normalized by average genome size (AGS), which was calculated using MicrobeCensus from each MGX samples^105^. Coriobacteriia <u>EA</u>-<u>m</u>etabolizing (*eam*) gene cluster abundance was calculated as an average of the gene abundance of *eah*, *eadh1*, and *eadh2*.

### Validation of urolithin A in metabolomics datasets

To confirm the presence of urolithin A in stool metabolomics from the PRISM IBD study, a synthetic standard was analyzed alongside reference pool material from the PRISM study using the same LC–MS profiling method (HILIC-neg) employed in the study^71^. Briefly, metabolites were extracted from PRISM stool homogenate (30 μL) with four volumes of 80% methanol containing inosine-^15^N4, thymine-d4 and glycocholate-d4 internal standards (Cambridge Isotope Laboratories). The extract was centrifuged (10 min, 9,000 g, 4 °C), and the supernatant was injected alongside 2 ng of urolithin A onto a 150 × 2.0-mm Luna NH2 column (Phenomenex). The column was eluted at a flow rate of 400 μL/min with initial conditions of 10% mobile phase A (20 mM ammonium acetate and 20 mM ammonium hydroxide in water) and 90% mobile phase B (10 mM ammonium hydroxide in 75:25 v/v acetonitrile/methanol) followed by a 10-min linear gradient to 100% mobile phase A. MS analyses were carried out using electrospray ionization in the negative ion mode using full scan analysis over *m/z* 60–750 at 70,000 resolution and 3 Hz data acquisition rate. Additional MS settings are: ion spray voltage, −3.0 kV; capillary temperature, 350 °C; probe heater temperature, 325 °C; sheath gas, 55; auxiliary gas, 10; and S-lens RF level 40. Product ion spectra MS/MS was generated with the same parameters at five different collision energies (HCD: 10, 20, 30, 40, 50 eV), and the *m/z* and retention time of the urolithin A reference standard were used to confirm the identity of the unknown feature in the PRISM data (Study Compound ID: HILn_PRISM_m12047, *m/z*: 227.0348, retention time: 2.78 min).

### Multivariate linear regression analysis

A ‘statsmodels’ python package (v0.14.1) was used to perform multivariate linear regression analysis between the gene abundances and urolithin A levels in paired metagenome and metabolome samples. Zero values were corrected by adding half the smallest non-zero value of each gene or metabolite level. To stabilize variance, abundances were log-transformed. For each IBD phenotype, we modelled the transformed abundance of urolithin A as a function of the *eam* gene cluster and *ucdh* abundances with age as a continuous covariate and four medications (antibiotics, immunosuppressants, mesalamine, and steroids) as binary covariates (ordinary least squares model). Nominal P values were adjusted for multiple hypothesis testing with a target FDR of 0.05 using Benjamini-Hochberg correction.

---

**Figure S1.**
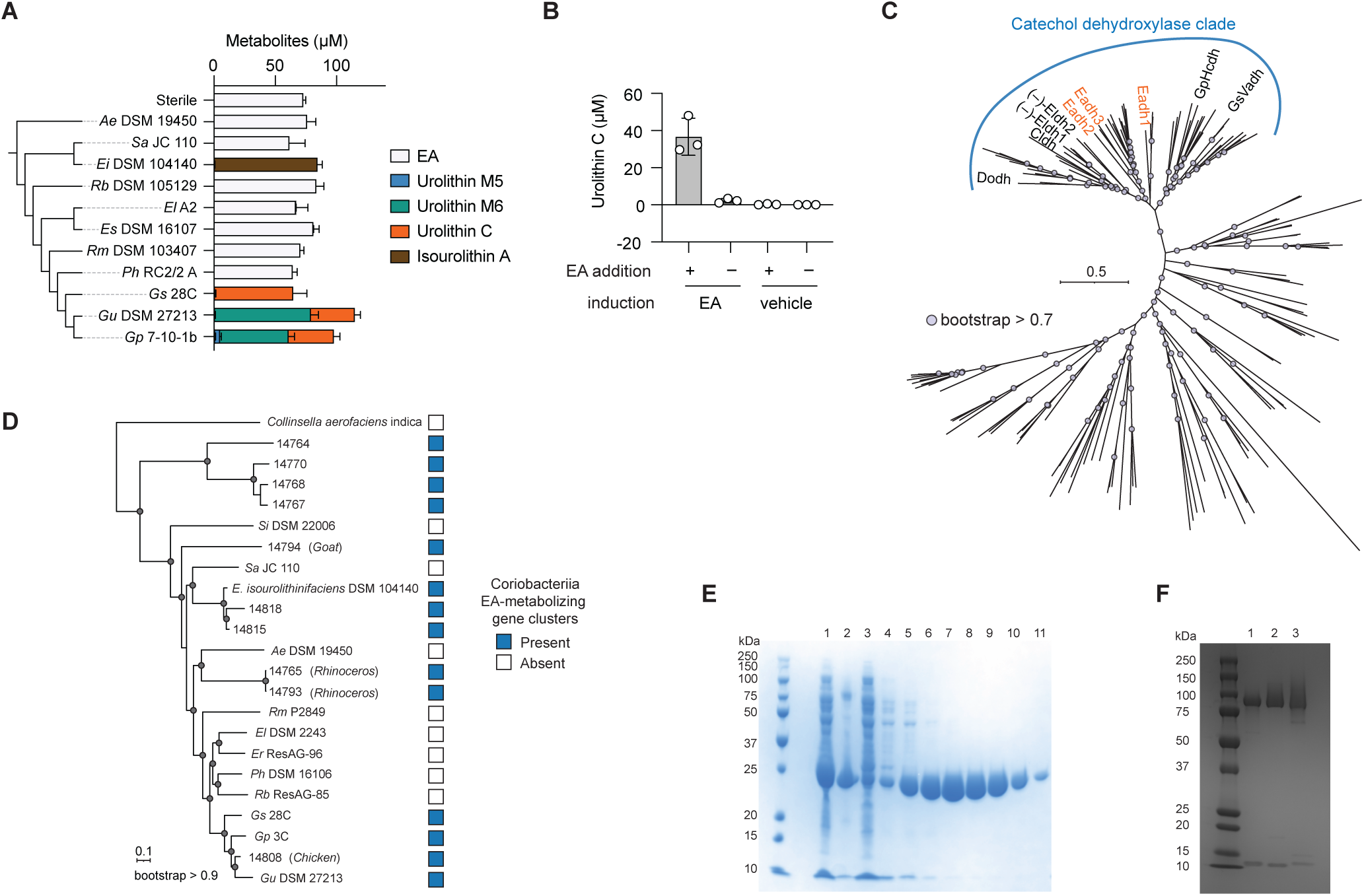
Identification of a Coriobacteriia gene cluster responsible for EA metabolism. **(A)** Endpoint concentrations of EA and urolithins in liquid cultures of Coriobacteriia strains anaerobically grown in medium containing 100 µM EA at 37 °C for 72 h. **(B)** Cell lysates of *Gs* 28C induced by 100 µM EA or vehicle were incubated with either 100 µM EA or vehicle at rt for 24 h. **(A–B)** Data represented as mean ± SD with n = 3 biological replicates. **(C)** Maximum-likelihood phylogenetic tree of molybdenum-dependent oxidoreductases encoded by *Gp* 3C, *Gp* 7-10-1b, *Gu* DSM 27213, *Gs* 28C, and *Ei* DSM 104140. Previously reported catechol dehydroxylases are labelled. Grey nodes indicate bootstrap value > 0.7. **(D)** Phylogenetic distribution of the *eam* gene cluster among animal gut-associated Coriobacteriia. Grey nodes indicate bootstrap value > 0.9. **(E)** SDS-PAGE image of fractions from Eah protein expression and purification. Lane annotations: 1, lysate; 2, pellet; 3, flow-through; 4–6, wash; 7–11, elution. **(F)** SDS-PAGE image of elution fractions of Eadh1 (lane 1), Eadh2 (lane 2), and Eadh3 (lane 3). *Ae, Adlercreutzia equolifaciens*; *Sa, Senegalimassilia anaerobia*; *Ei, Ellagibacter isourolithinifaciens*; *Rb, Rubneribacter badeniensis*; *El, Eggerthella lenta*; *Es, Eggerthella sinensis*; *Rm, Raoultibacter massiliensis*; *Ph, Paraeggerthella hongkongensis*; *Gs, Gordonibacter species*; *Gu, Gordonibacter urolithinfaciens*; *Gp, Gordonibacter pamelaeae*; *Si, Slackia isoflavoniconvertens*; *Er, Enteroscipio rubneri*; Dodh, DOPAC dehydroxylase; Cldh, catechol lignan dehydroxylase; (–)-Eldh1, (–)- dihydroxyenterolactone dehydroxylase 1; (–)-Eldh2, (–)-dihydroxyenterolactone dehydroxylase 2; *Gp* Hcdh, *Gordonibacter pamelaeae* hydrocaffeic acid dehydroxylase; *Gs* Vadh, *Gs* 28C 5-(3’,4’-dihydroxyphenyl)valeric acid dehydroxylase.

**Figure S2.**
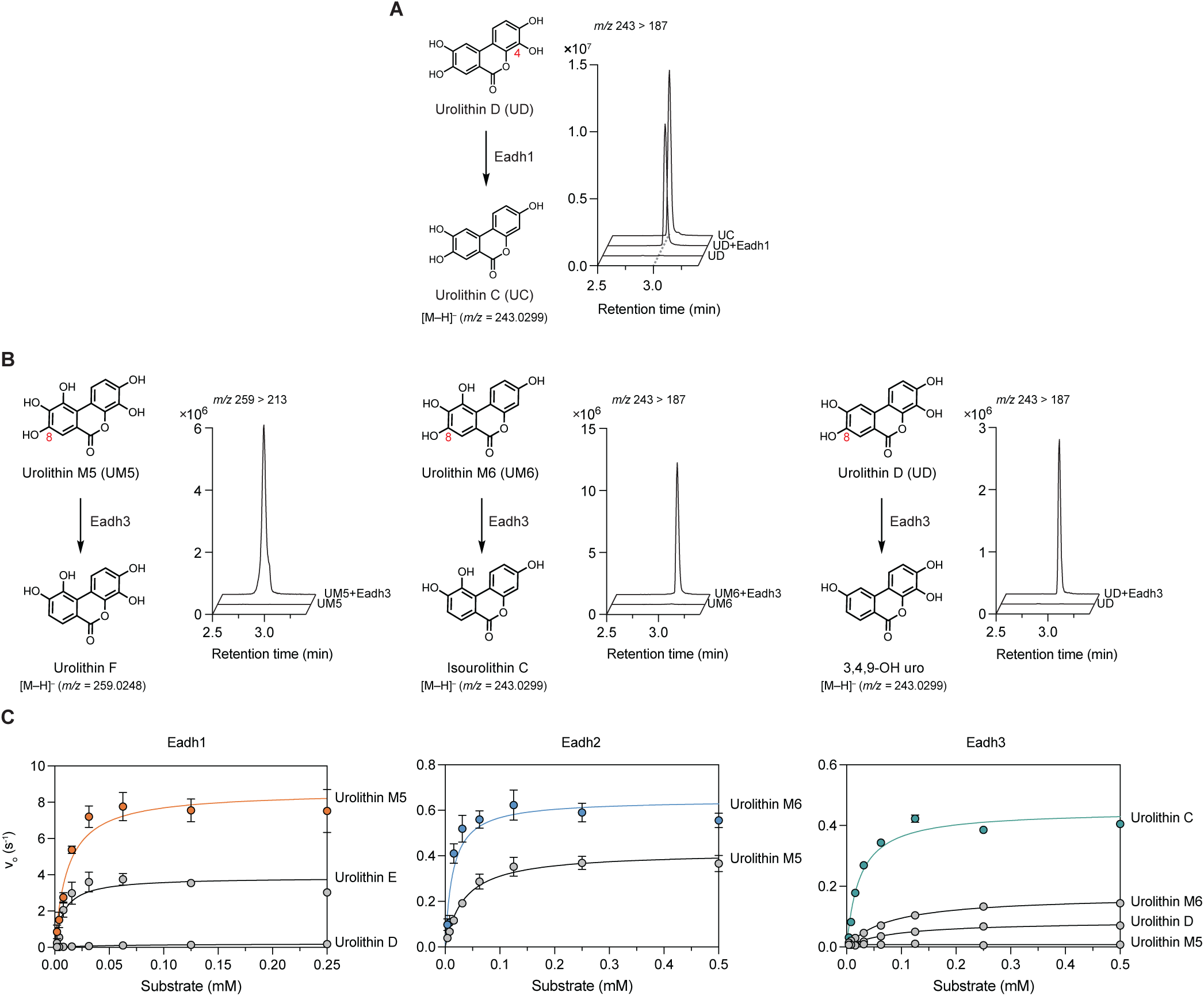
Catechol dehydroxylase substrate specificity and kinetics explain urolithin production. **(A)** LC–MS/MS traces of urolithin D dehydroxylation by Eadh1. **(B)** LC–MS/MS traces of urolithin metabolism by Eadh3 toward urolithins containing C8-OH. Structures of predicted products are shown. **(A–B)** 100 nM enzyme was incubated with 100 µM substrate, 200 µM methyl viologen, and 100 µM sodium dithionite at rt for 20 h in a pH 7.0 buffer (20 mM MOPS and 300 mM NaCl) under anaerobic conditions. **(C)** Michaelis–Menten curves of Eadh1, Eadh2, and Eadh3 towards various substrates. The reactions were performed anaerobically in a pH 7.0 buffer (20 mM MOPS and 300 mM NaCl) containing varying concentrations of enzymes and substrates, 200 µM methyl viologen, and 100 µM sodium dithionite at 30 °C.

**Figure S3.**
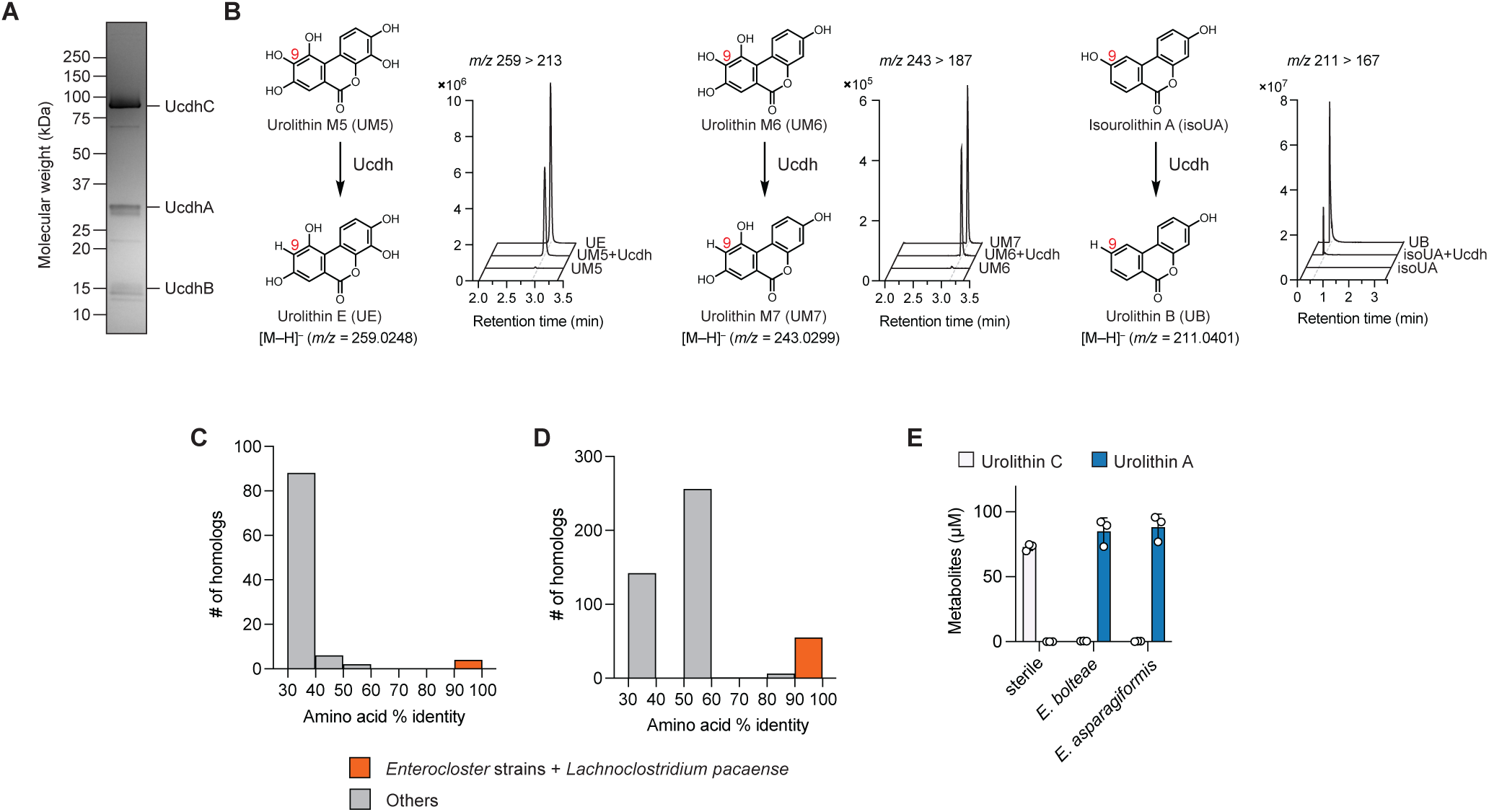
A previously unappreciated xanthine oxidase homolog catalyzes dehydroxylation of urolithin C to urolithin A. **(A)** SDS-PAGE image of an elution fraction of Ucdh. **(B)** LC–MS/MS traces of dehydroxylated products after C9-OH containing urolithins were incubated with Ucdh. 2 µM Ucdh was incubated with 100 µM substrate and 1 mM NADH at rt for 20 h in a pH 7.0 buffer (20 mM MOPS and 300 mM NaCl) under anaerobic conditions. **(C)** A histogram listing top 100 hits with highest amino acid percent identity to Ucdh from the NCBI RefSeq Select protein database. **(D)** A histogram listing Ucdh homologs found in a human-associated microbial assembled genome dataset. **(E)** Urolithin C dehydroxylation to urolithin A by Ucdh-encoding species. Cultures were grown with 100 µM urolithin C at 37 °C for 72 h (n=3 biological triplicates).

**Figure S4.**
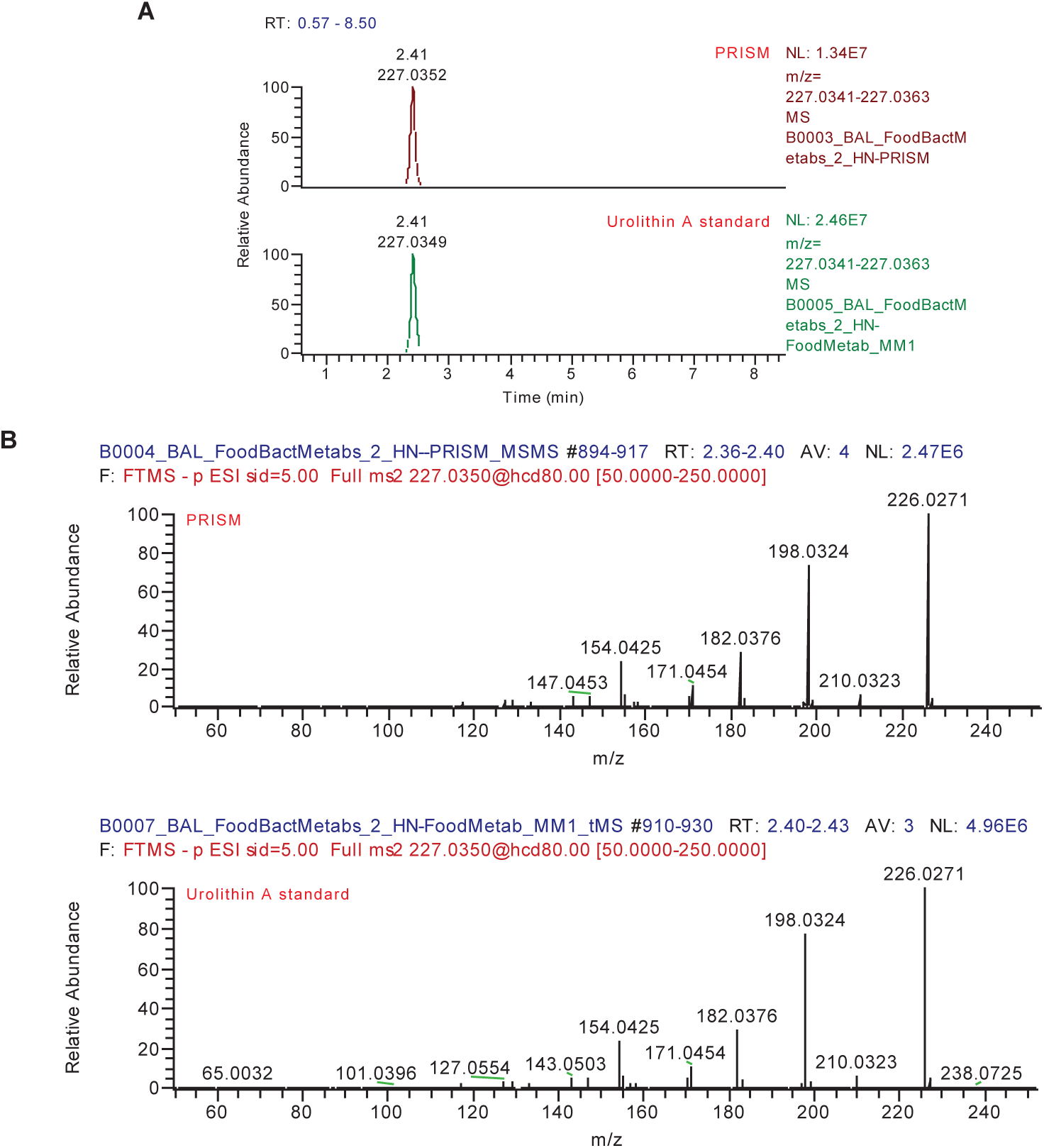
Identification of urolithin A in PRISM metabolome database. **(A)** Extracted ion chromatogram (EIC) of urolithin A standard (bottom) and a corresponding mass peak (QI1103) in PRISM metabolome database (top). **(B)** MS/MS spectra of urolithin A standard (bottom) and QI1103 (top).

**Figure S5.**
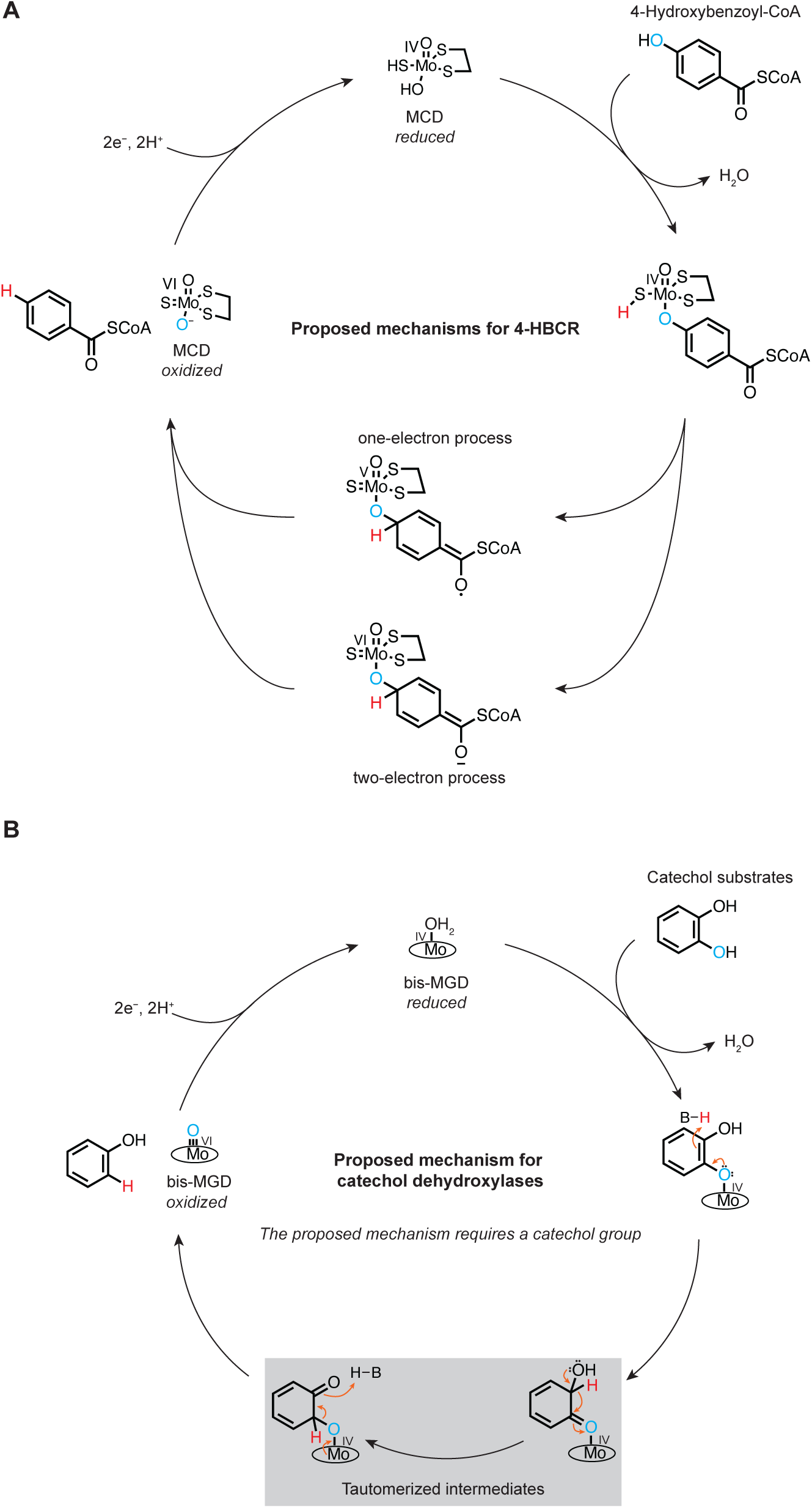
Proposed mechanisms for 4-HBCR and catechol dehydroxylases. **(A)** Two proposed mechanisms for 4-HBCR. **(B)** Proposed mechanism for catechol dehydroxylases via tautomerization.

**Table S1.**
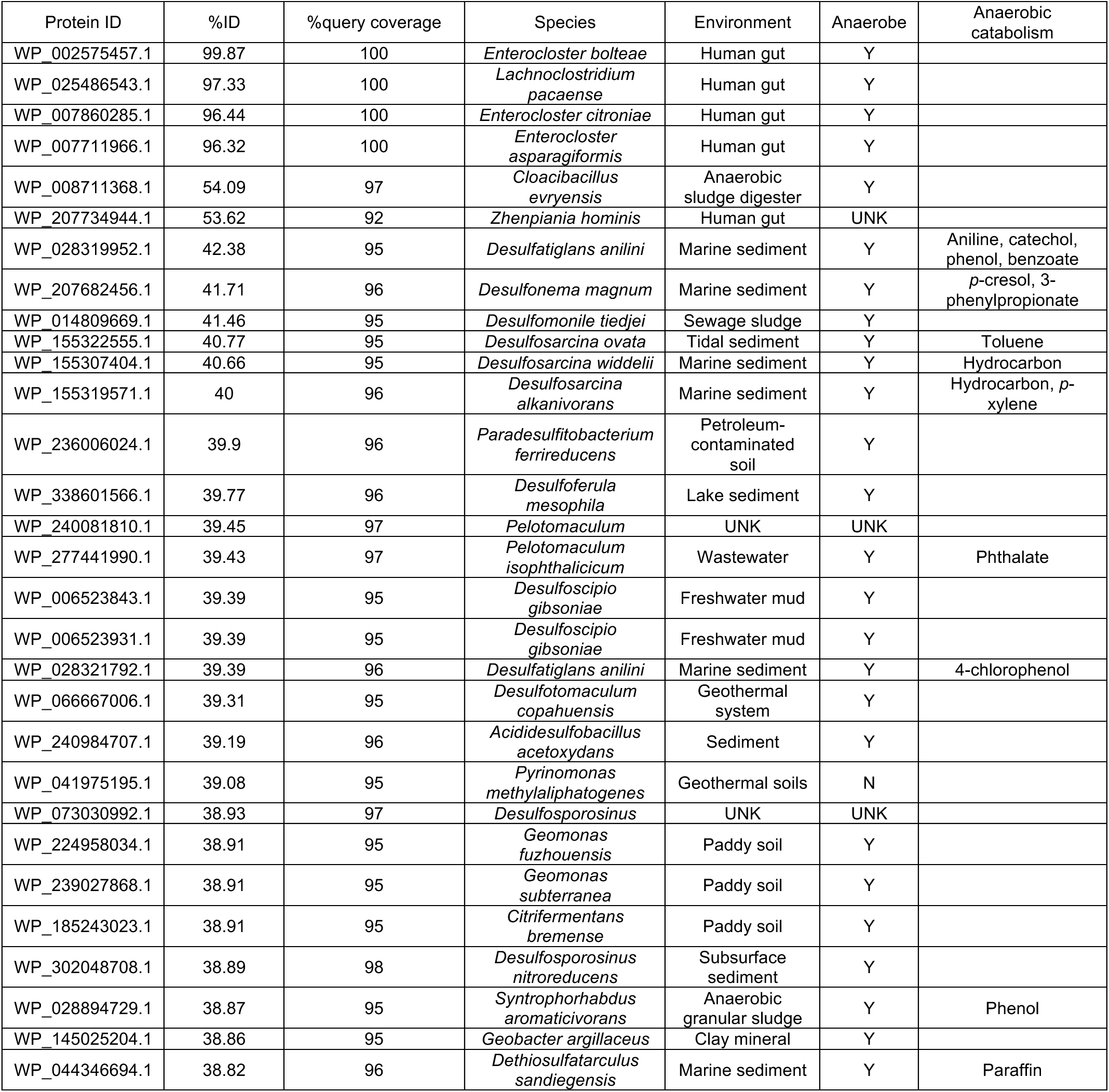
Top 30 highest homology hits of Ucdh among the NCBI RefSeq Select protein database and their encoding species. UNK, unknown.

## Reference

1. García-Villalba, R. et al. Urolithins: a Comprehensive Update on their Metabolism, Bioactivity, and Associated Gut Microbiota. Molecular Nutrition & Food Research 66, 2101019 (2022).

2. Bao, Y. et al. Association of Nut Consumption with Total and Cause-Specific Mortality. New England Journal of Medicine 369, 2001–2011 (2013).

3. Basu, A., Rhone, M. & Lyons, T. J. Berries: emerging impact on cardiovascular health. Nutrition Reviews 68, 168–177 (2010).

4. Yang, B. & Kortesniemi, M. Clinical evidence on potential health benefits of berries. Current Opinion in Food Science 2, 36–42 (2015).

5. Guasch, -Ferré Marta et al. Nut Consumption and Risk of Cardiovascular Disease. Journal of the American College of Cardiology 70, 2519–2532 (2017).

6. Doyle, B. & Griffiths, L. A. The metabolism of ellagic acid in the rat. Xenobiotica 10, 247–256 (1980).

7. Ryu, D. et al. Urolithin A induces mitophagy and prolongs lifespan in C. elegans and increases muscle function in rodents. Nature Medicine 22, 879–888 (2016).

8. Ghosh, N. et al. Urolithin A augments angiogenic pathways in skeletal muscle by bolstering NAD + and SIRT1. Scientific Reports 10, 20184 (2020).

9. Abdelazeem, K. N. M., Kalo, M. Z., Beer-Hammer, S. & Lang, F. The gut microbiota metabolite urolithin A inhibits NF-κB activation in LPS stimulated BMDMs. Scientific Reports 11, 7117 (2021).

10. Cho, S. I., Jo, E.-R. & Song, H. Urolithin A attenuates auditory cell senescence by activating mitophagy. Sci Rep 12, 7704 (2022).

11. Jiménez-Loygorri, J. I. et al. Mitophagy curtails cytosolic mtDNA-dependent activation of cGAS/STING inflammation during aging. Nat Commun 15, 830 (2024).

12. Singh, R. et al. Enhancement of the gut barrier integrity by a microbial metabolite through the Nrf2 pathway. Nature Communications 10, 1–18 (2019).

13. Toney, A. M., Fox, D., Chaidez, V., Ramer-Tait, A. E. & Chung, S. Immunomodulatory Role of Urolithin A on Metabolic Diseases. Biomedicines 9, 192 (2021).

14. Ma, M. et al. Urolithin A Alleviates Colitis in Mice by Improving Gut Microbiota Dysbiosis, Modulating Microbial Tryptophan Metabolism, and Triggering AhR Activation. J. Agric. Food Chem. (2023) doi:10.1021/acs.jafc.3c00830.

15. Denk, D. et al. Expansion of T memory stem cells with superior anti-tumor immunity by Urolithin A-induced mitophagy. Immunity 55, 2059–2073.e8 (2022).

16. Girotra, M. et al. Induction of mitochondrial recycling reverts age-associated decline of the hematopoietic and immune systems. Nat Aging 1–10 (2023) doi:10.1038/s43587-023-00473-3.

17. Luan, P. et al. Urolithin A improves muscle function by inducing mitophagy in muscular dystrophy. Science Translational Medicine 13, (2021).

18. Fang, E. F. et al. Mitophagy inhibits amyloid-β and tau pathology and reverses cognitive deficits in models of Alzheimer’s disease. Nat Neurosci 22, 401–412 (2019).

19. Gong, Z. et al. Urolithin A attenuates memory impairment and neuroinflammation in APP/PS1 mice. J Neuroinflammation 16, 62 (2019).

20. Shen, P.-X. et al. Urolithin A ameliorates experimental autoimmune encephalomyelitis by targeting aryl hydrocarbon receptor. eBioMedicine 64, (2021).

21. Toney, A. M. et al. Urolithin A, a Gut Metabolite, Improves Insulin Sensitivity Through Augmentation of Mitochondrial Function and Biogenesis. Obesity 27, 612–620 (2019).

22. Xia, B. et al. Urolithin A exerts antiobesity effects through enhancing adipose tissue thermogenesis in mice. PLOS Biology 18, e3000688 (2020).

23. Yang, J. et al. Ellagic Acid and Its Microbial Metabolite Urolithin A Alleviate Diet-Induced Insulin Resistance in Mice. Molecular Nutrition & Food Research 64, 2000091 (2020).

24. Andreux, P. A. et al. The mitophagy activator urolithin A is safe and induces a molecular signature of improved mitochondrial and cellular health in humans. Nat Metab 1, 595–603 (2019).

25. Singh, A. et al. Urolithin A improves muscle strength, exercise performance, and biomarkers of mitochondrial health in a randomized trial in middle-aged adults. Cell Reports Medicine 3, 100633 (2022).

26. Liu, S. et al. Effect of Urolithin A Supplementation on Muscle Endurance and Mitochondrial Health in Older Adults: A Randomized Clinical Trial. JAMA Network Open 5, e2144279 (2022).

27. Tomás-Barberán, F. A., García-Villalba, R., González-Sarrías, A., Selma, M. V. & Espín, J. C. Ellagic Acid Metabolism by Human Gut Microbiota: Consistent Observation of Three Urolithin Phenotypes in Intervention Trials, Independent of Food Source, Age, and Health Status. J. Agric. Food Chem. 62, 6535–6538 (2014).

28. Romo-Vaquero, M. et al. Interindividual variability in the human metabolism of ellagic acid: Contribution of Gordonibacter to urolithin production. Journal of Functional Foods 17, 785–791 (2015).

29. Cortés-Martín, A. et al. The gut microbiota urolithin metabotypes revisited: the human metabolism of ellagic acid is mainly determined by aging. Food Funct. 9, 4100–4106 (2018).

30. Selma, M. V., Beltrán, D., García-Villalba, R., Espín, J. C. & Tomás-Barberán, F. A. Description of urolithin production capacity from ellagic acid of two human intestinal Gordonibacter species. Food Funct. 5, 1779–1784 (2014).

31. Iglesias-Aguirre, C. E. et al. Gut Bacteria Involved in Ellagic Acid Metabolism To Yield Human Urolithin Metabotypes Revealed. J. Agric. Food Chem. (2023) doi:10.1021/acs.jafc.2c08889.

32. Selma, M. V. et al. Isolation of Human Intestinal Bacteria Capable of Producing the Bioactive Metabolite Isourolithin A from Ellagic Acid. Front. Microbiol. 8, (2017).

33. Kang, I., Kim, Y., Tomás-Barberán, F. A., Espín, J. C. & Chung, S. Urolithin A, C, and D, but not iso-urolithin A and urolithin B, attenuate triglyceride accumulation in human cultures of adipocytes and hepatocytes. Molecular Nutrition & Food Research 60, 1129–1138 (2016).

34. Giménez-Bastida, J. A., Ávila-Gálvez, M. Á., Espín, J. C. & González-Sarrías, A. The gut microbiota metabolite urolithin A, but not other relevant urolithins, induces p53-dependent cellular senescence in human colon cancer cells. Food and Chemical Toxicology 139, 111260 (2020).

35. González-Sarrías, A., Núñez-Sánchez, M. Á., García-Villalba, R., Tomás-Barberán, F. A. & Espín, J. C. Antiproliferative activity of the ellagic acid-derived gut microbiota isourolithin A and comparison with its urolithin A isomer: the role of cell metabolism. Eur J Nutr 56, 831–841 (2017).

36. Selma, M. V. et al. The human gut microbial ecology associated with overweight and obesity determines ellagic acid metabolism. Food Funct. 7, 1769–1774 (2016).

37. Selma, M. V. et al. The gut microbiota metabolism of pomegranate or walnut ellagitannins yields two urolithin-metabotypes that correlate with cardiometabolic risk biomarkers: Comparison between normoweight, overweight-obesity and metabolic syndrome. Clinical Nutrition 37, 897–905 (2018).

38. Cortés-Martín, A., Colmenarejo, G., Selma, M. V. & Espín, J. C. Genetic Polymorphisms, Mediterranean Diet and Microbiota-Associated Urolithin Metabotypes can Predict Obesity in Childhood-Adolescence. Sci Rep 10, 7850 (2020).

39. Li, C., Zhao, X., Wang, A., Huber, G. W. & Zhang, T. Catalytic Transformation of Lignin for the Production of Chemicals and Fuels. Chem. Rev. 115, 11559–11624 (2015).

40. Saidi, M. et al. Upgrading of lignin-derived bio-oils by catalytic hydrodeoxygenation. Energy Environ. Sci. 7, 103–129 (2013).

41. Duan, H. et al. Molecular nitrogen promotes catalytic hydrodeoxygenation. Nat Catal 2, 1078–1087 (2019).

42. Maini Rekdal, V., Bess, E. N., Bisanz, J. E., Turnbaugh, P. J. & Balskus, E. P. Discovery and inhibition of an interspecies gut bacterial pathway for Levodopa metabolism. Science 364, eaau6323 (2019).

43. Bess, E. N. et al. Genetic basis for the cooperative bioactivation of plant lignans by Eggerthella lenta and other human gut bacteria. Nature Microbiology 5, 56–66 (2020).

44. Maini Rekdal, V., et al. A widely distributed metalloenzyme class enables gut microbial metabolism of host-and diet-derived catechols. eLife 9, e50845 (2020).

45. Le, C. (Chip), Bae, M., Kiamehr, S. & Balskus, E. P. Emerging Chemical Diversity and Potential Applications of Enzymes in the DMSO Reductase Superfamily. Annual Review of Biochemistry 91, 475–504 (2022).

46. Zahn, L. E., Gannon, P. M. & Rajakovich, L. J. Iron-sulfur cluster-dependent enzymes and molybdenum-dependent reductases in the anaerobic metabolism of human gut microbes. Metallomics 16, mfae049 (2024).

47. Bae, M. et al. Metatranscriptomics-guided discovery and characterization of a polyphenol-metabolizing gut microbial enzyme. Cell Host & Microbe 1–10 doi:10.1016/j.chom.2024.10.002.

48. Dong, X., Bae, M., Le, C. (Chip), Aguilar Ramos, M. A. & Balskus, E. P. Enantiocomplementary gut bacterial enzymes metabolize dietary polyphenols. in revision.

49. Little, A. S. et al. Dietary- and host-derived metabolites are used by diverse gut bacteria for anaerobic respiration. Nat Microbiol 9, 55–69 (2024).

50. Carmona, M. et al. Anaerobic Catabolism of Aromatic Compounds: a Genetic and Genomic View. Microbiol. Mol. Biol. Rev. 73, 71–133 (2009).

51. Gibson, J., Dispensa, M. & Harwood, C. S. 4-hydroxybenzoyl coenzyme A reductase (dehydroxylating) is required for anaerobic degradation of 4-hydroxybenzoate by Rhodopseudomonas palustris and shares features with molybdenum-containing hydroxylases. Journal of Bacteriology 179, 634–642 (1997).

52. Unciuleac, M., Warkentin, E., Page, C. C., Boll, M. & Ermler, U. Structure of a Xanthine Oxidase-Related 4-Hydroxybenzoyl-CoA Reductase with an Additional [4Fe-4S] Cluster and an Inverted Electron Flow. Structure 12, 2249–2256 (2004).

53. Dong, X. et al. Genetic manipulation of the human gut bacterium Eggerthella lenta reveals a widespread family of transcriptional regulators. Nat Commun 13, 7624 (2022).

54. Altschul, S. F., Gish, W., Miller, W., Myers, E. W. & Lipman, D. J. Basic local alignment search tool. Journal of Molecular Biology 215, 403–410 (1990).

55. Paysan-Lafosse, T. et al. InterPro in 2022. Nucleic Acids Research 51, D418–D427 (2023).

56. Espín, J. C. et al. Iberian Pig as a Model To Clarify Obscure Points in the Bioavailability and Metabolism of Ellagitannins in Humans. J. Agric. Food Chem. 55, 10476–10485 (2007).

57. Boll, M. et al. Redox Centers of 4-Hydroxybenzoyl-CoA Reductase, a Member of the Xanthine Oxidase Family of Molybdenum-containing Enzymes *. Journal of Biological Chemistry 276, 47853–47862 (2001).

58. El Kasmi, A., Brachmann, R., Fuchs, G. & Ragsdale, S. W. Hydroxybenzoyl-CoA reductase: coupling kinetics and electrochemistry to derive enzyme mechanisms. Biochemistry 34, 11668–11677 (1995).

59. Neumann, M., Schulte, M., Jünemann, N., Stöcklein, W. & Leimkühler, S. Rhodobacter capsulatus XdhC Is Involved in Molybdenum Cofactor Binding and Insertion into Xanthine Dehydrogenase *. Journal of Biological Chemistry 281, 15701–15708 (2006).

60. Breese, K. & Fuchs, G. 4-Hydroxybenzoyl-CoA reductase (dehydroxylating) from the denitrifying bacterium Thauera aromatica. European Journal of Biochemistry 251, 916–923 (1998).

61. Palmer, T. et al. Involvement of the narJ and mob gene products in distinct steps in the biosynthesis of the molybdoenzyme nitrate reductase in Escherichia coli. Molecular Microbiology 20, 875–884 (1996).

62. Hitch, T. C. A. et al. Broad diversity of human gut bacteria accessible via a traceable strain deposition system. 2024.06.20.599854 Preprint at 10.1101/2024.06.20.599854 (2024).

63. Hosny, M. et al. Clostridium pacaense:a new species within the genus Clostridium. New Microbes and New Infections 28, 6–10 (2019).

64. Davidova, I. A. et al. Dethiosulfatarculus sandiegensis gen. nov., sp. nov., isolated from a methanogenic paraffin-degrading enrichment culture and emended description of the family Desulfarculaceae. International Journal of Systematic and Evolutionary Microbiology 66, 1242–1248 (2016).

65. Suzuki, D., Li, Z., Cui, X., Zhang, C. & Katayama, A. Reclassification of Desulfobacterium anilini as Desulfatiglans anilini comb. nov. within Desulfatiglans gen. nov.w, and description of a 4-chlorophenol-degrading sulfate-reducing bacterium, Desulfatiglans parachlorophenolica sp. nov. International Journal of Systematic and Evolutionary Microbiology 64, 3081–3086 (2014).

66. Qiu, Y.-L. et al. Pelotomaculum terephthalicum sp. nov. and Pelotomaculum isophthalicum sp. nov.: two anaerobic bacteria that degrade phthalate isomers in syntrophic association with hydrogenotrophic methanogens. Arch Microbiol 185, 172–182 (2006).

67. Watanabe, M., Higashioka, Y., Kojima, H. & Fukui, M. Desulfosarcina widdelii sp. nov. and Desulfosarcina alkanivorans sp. nov., hydrocarbon-degrading sulfate-reducing bacteria isolated from marine sediment and emended description of the genus Desulfosarcina. International Journal of Systematic and Evolutionary Microbiology 67, 2994–2997 (2017).

68. Watanabe, M., Higashioka, Y., Kojima, H. & Fukui, M. Proposal of *Desulfosarcina ovata* subsp. *sediminis* subsp. nov., a novel toluene-degrading sulfate-reducing bacterium isolated from tidal flat sediment of Tokyo Bay. Systematic and Applied Microbiology 43, 126109 (2020).

69. Schnaars, V. et al. Proteogenomic Insights into the Physiology of Marine, Sulfate-Reducing, Filamentous Desulfonema limicola and Desulfonema magnum. Microbial Physiology 31, 36–56 (2021).

70. Schnell, S., Bak, F. & Pfennig, N. Anaerobic degradation of aniline and dihydroxybenzenes by newly isolated sulfate-reducing bacteria and description of Desulfobacterium anilini. Arch. Microbiol. 152, 556–563 (1989).

71. Franzosa, E. A. et al. Gut microbiome structure and metabolic activity in inflammatory bowel disease. Nat Microbiol 4, 293–305 (2019).

72. Lloyd-Price, J. et al. Multi-omics of the gut microbial ecosystem in inflammatory bowel diseases. Nature 569, 655–662 (2019).

73. Vila, A. V. et al. Faecal metabolome and its determinants in inflammatory bowel disease. Gut 72, 1472–1485 (2023).

74. Ning, L. et al. Microbiome and metabolome features in inflammatory bowel disease via multi-omics integration analyses across cohorts. Nat Commun 14, 7135 (2023).

75. Lu, Z. & Imlay, J. A. When anaerobes encounter oxygen: mechanisms of oxygen toxicity, tolerance and defence. Nat Rev Microbiol 19, 774–785 (2021).

76. Rigottier-Gois, L. Dysbiosis in inflammatory bowel diseases: the oxygen hypothesis. ISME J 7, 1256–1261 (2013).

77. Hille, R., Hall, J. & Basu, P. The Mononuclear Molybdenum Enzymes. Chem. Rev. 114, 3963–4038 (2014).

78. Yang, J. et al. Active Site Structure and Mechanism of a Molybdenum Catechol Dehydroxylase. Submitted.

79. García-Villalba, R., Selma, M. V., Espín, J. C. & Tomás-Barberán, F. A. Identification of Novel Urolithin Metabolites in Human Feces and Urine after the Intake of a Pomegranate Extract. J. Agric. Food Chem. 67, 11099–11107 (2019).

80. Gaya, P., Medina, M., Sánchez-Jiménez, A. & Landete, J. M. Phytoestrogen Metabolism by Adult Human Gut Microbiota. Molecules 21, 1034 (2016).

81. Pidgeon, R. et al. The dietary ellagitannin metabolite urolithin A is produced by a molybdenum-dependent dehydroxylase encoded by prevalent human gut Enterocloster spp. 2024.02.08.579493 Preprint at 10.1101/2024.02.08.579493 (2024).

82. Liu, Y. et al. A widely distributed gene cluster compensates for uricase loss in hominids. Cell 186, 3400–3413.e20 (2023).

83. Feng, C. et al. Discovery of a Gut Bacterial Pathway for Ergothioneine Catabolism. J. Am. Chem. Soc. (2024) doi:10.1021/jacs.4c09350.

84. Zhang, X. et al. Isolation and characterization of a novel human intestinal Enterococcus faecium FUA027 capable of producing urolithin A from ellagic acid. Frontiers in Nutrition 9, (2022).

85. Mi, H. et al. Lactococcus garvieae FUA009, a Novel Intestinal Bacterium Capable of Producing the Bioactive Metabolite Urolithin A from Ellagic Acid. Foods 11, 2621 (2022).

86. Gaya, P., Peirotén, Á., Medina, M., Álvarez, I. & Landete, J. M. Bifidobacterium pseudocatenulatum INIA P815: The first bacterium able to produce urolithins A and B from ellagic acid. Journal of Functional Foods 45, 95–99 (2018).

87. Shishkin, A. A. et al. Simultaneous generation of many RNA-seq libraries in a single reaction. Nat Methods 12, 323–325 (2015).

88. Langmead, B. & Salzberg, S. L. Fast gapped-read alignment with Bowtie 2. Nat Methods 9, 357–359 (2012).

89. Putri, G. H., Anders, S., Pyl, P. T., Pimanda, J. E. & Zanini, F. Analysing high-throughput sequencing data in Python with HTSeq 2.0. Bioinformatics 38, 2943–2945 (2022).

90. Love, M. I., Huber, W. & Anders, S. Moderated estimation of fold change and dispersion for RNA-seq data with DESeq2. Genome Biology 15, 550 (2014).

91. Zhu, A., Ibrahim, J. G. & Love, M. I. Heavy-tailed prior distributions for sequence count data: removing the noise and preserving large differences. Bioinformatics 35, 2084–2092 (2019).

92. Ritchie, M. E. et al. limma powers differential expression analyses for RNA-sequencing and microarray studies. Nucleic Acids Research 43, e47 (2015).

93. Fu, L., Niu, B., Zhu, Z., Wu, S. & Li, W. CD-HIT: accelerated for clustering the next-generation sequencing data. Bioinformatics 28, 3150–3152 (2012).

94. Katoh, K. & Standley, D. M. MAFFT Multiple Sequence Alignment Software Version 7: Improvements in Performance and Usability. Molecular Biology and Evolution 30, 772–780 (2013).

95. Capella-Gutiérrez, S., Silla-Martínez, J. M. & Gabaldón, T. trimAl: a tool for automated alignment trimming in large-scale phylogenetic analyses. Bioinformatics 25, 1972–1973 (2009).

96. Minh, B. Q. et al. IQ-TREE 2: New Models and Efficient Methods for Phylogenetic Inference in the Genomic Era. Molecular Biology and Evolution 37, 1530–1534 (2020).

97. Pasolli, E. et al. Extensive Unexplored Human Microbiome Diversity Revealed by Over 150,000 Genomes from Metagenomes Spanning Age, Geography, and Lifestyle. Cell 176, 649–662.e20 (2019).

98. Stewart, R. D. et al. Compendium of 4,941 rumen metagenome-assembled genomes for rumen microbiome biology and enzyme discovery. Nature Biotechnology 37, 953–961 (2019).

99. Wilkinson, T. et al. 1200 high-quality metagenome-assembled genomes from the rumen of African cattle and their relevance in the context of sub-optimal feeding. Genome Biology 21, 229 (2020).

100. Glendinning, L., Stewart, R. D., Pallen, M. J., Watson, K. A. & Watson, M. Assembly of hundreds of novel bacterial genomes from the chicken caecum. Genome Biology 21, 34 (2020).

101. Youngblut, N. D. et al. Large-Scale Metagenome Assembly Reveals Novel Animal-Associated Microbial Genomes, Biosynthetic Gene Clusters, and Other Genetic Diversity. mSystems 5, 10.1128/msystems.01045-20 (2020).

102. Nayfach, S. et al. A genomic catalog of Earth’s microbiomes. Nat Biotechnol 39, 499–509 (2021).

103. Arkin, A. P. et al. KBase: The United States Department of Energy Systems Biology Knowledgebase. Nat Biotechnol 36, 566–569 (2018).

104. Buchfink, B., Xie, C. & Huson, D. H. Fast and sensitive protein alignment using DIAMOND. Nat Methods 12, 59–60 (2015).

105. Nayfach, S. & Pollard, K. S. Average genome size estimation improves comparative metagenomics and sheds light on the functional ecology of the human microbiome. Genome Biology 16, 51 (2015).

